# Hepatitis B virus protein X promotes hepatocyte plasticity and survival in a differentiated human liver organoid system

**DOI:** 10.64898/2026.06.26.734750

**Authors:** Xingyu Fan, Bram Torenvliet, Alexandros Galaras, Tanvir Hossain, Lincon Hasda, Martin E van Royen, Helmuth Gehart, Lili Zhao, Eleni Katsoni, Tsung Wai Kan, Panagiotis Moulos, Shringar Rao, Farzin Pourfarzad, Javier Frias Aldeguer, Sylvia F. Boj, Pantelis Hatzis, Robert-Jan Palstra, Tokameh Mahmoudi

**Author notes:** **Corresponding author:** Tokameh Mahmoudi, Address: Departments of Pathology and Urology, Erasmus University Medical Center, Netherlands, Email address. These authors contributed equally (Xingyu Fan, Bram Torenvliet).

## Abstract

**Background & Aims:** Hepatitis B virus (HBV) drives hepatocellular carcinoma in part through the activity of its X protein (HBx), yet the mechanisms by which HBx alters hepatocyte function remain incompletely understood. Progress has been limited by the lack of relevant human models that support controlled HBx expression in mature hepatocytes. Here, we use an improved hepatocyte-like organoid (HLO) platform that supports enhanced hepatocyte maturation to investigate HBx function in a differentiated hepatocyte context.

**Methods:** Adult stem cell-derived HLOs were differentiated using an optimized protocol to generate hepatocyte-like cells with enhanced maturation and transcriptional similarity to primary liver tissue. HBx function was interrogated using both cognate promoter-driven expression and doxycycline-inducible systems across multiple donor-derived organoid lines. Transcriptomic, pathway, and single-cell imaging analyses were performed to assess the impact of HBx expression on hepatocytes.

**Results:** HBx expression consistently suppressed apoptosis-associated transcripts and reduced expression of core hepatocyte identity genes, including CYP3A4. Pathway analysis revealed downregulation of liver-specific functions, including metabolism, detoxification, complement, and coagulation. At the single-cell level, higher HBx expression was associated with reduced caspase 3/7 activation following apoptotic challenge and decreased hepatocyte marker expression. Functionally, HBx expression increased resistance to apoptosis and enhanced the ability of differentiated hepatocyte-like cells to revert to a proliferative, less differentiated state.

**Conclusions:** HBx expression in differentiated human liver organoids reduces apoptosis and impairs hepatocyte identity, consistently across donors and expression systems. These findings support a model in which HBx promotes a survival-permissive less differentiated state that may contribute to early HBV-driven tumorigenesis. This HLO platform provides a relevant system to dissect HBV-host interactions and reveals a mechanism by which HBV may prime the liver for malignant transformation.

**Impact and implications:** Understanding how HBV promotes hepatocellular carcinoma remains a critical challenge, partly due to the lack of physiologically relevant human derived model systems to study HBx function. Using a differentiated adult human liver organoid system, we show that HBx simultaneously suppresses apoptosis and disrupts hepatocyte identity, providing a mechanistic framework for how HBV may prime hepatocytes for malignant transformation. These findings are particularly relevant for researchers studying HBV pathogenesis and liver cancer, as well as for clinicians aiming to better understand early disease progression. While further validation in more complex multicellular systems is needed, this platform can support the identification of HBx-targeted therapeutic strategies and guide the development of improved adult human derived models for virus-host interaction studies.

## Introduction

Hepatitis B virus (HBV) infection remains a major global health burden, with approximately 254 million people chronically infected despite the availability of effective vaccines and antiviral therapies (1). Chronic HBV infection is a leading risk factor for hepatocellular carcinoma (HCC), conferring up to a 30-fold increased risk, and accounts for more than 50% of worldwide liver cancer-related deaths (2, 3). HBV infection is thought to promote hepatocarcinogenesis through multiple mechanisms reflecting a complex interplay between persistent viral infection and host immune responses (4). Continuous immune-mediated inflammation in the liver leads to repeated cycles of hepatocyte injury, cell death, and compensatory proliferation as the tissue attempts to regenerate. This chronic regenerative pressure increases the likelihood of accumulating genetic and epigenetic alterations in hepatocytes. In addition to host-driven inflammatory processes, virus-specific mechanisms are thought to contribute to oncogenic transformation (5–7). For example, integration of fragments of the viral genome into the host hepatocyte genome can induce insertional mutagenesis and chromosomal instability, in turn resulting in dysregulation of host gene expression, and persistent oncogenic signalling (8). Dysregulation of host gene expression and signalling pathways can occur via viral promoters and enhancers (9). Expression of viral proteins, most notably the hepatitis B virus protein (HBx), is also thought to drive oncogenic processes through broad regulatory effects on hepatocytes (10, 11).

HBx is encoded by the smallest open reading frame within the highly compact 3.2 kb HBV genome (12). In the context of the viral life cycle, HBx enhances transcription from covalently closed circular DNA (cccDNA) and modulates host pathways required for viral replication and persistence (13). Notably, the HBx coding region is among the most frequently integrated regions of the HBV genome, enabling sustained HBx expression even in the absence of active viral replication (14, 15). Mechanistically, HBx is thought to exert its effects primarily through interactions with host proteins, leading to dysregulation of key signalling pathways, inhibition of p53 activity, disruption of cell cycle control, and impaired DNA repair (16–20).

Despite extensive study, the molecular mechanisms underlying HBx function in hepatocarcinogenesis remain incompletely understood, in part due to limitations of existing experimental models. Reported effects of HBx vary substantially depending on model system, experimental design and HBx expression level (21, 22). Immortalized hepatoma cell lines, which are widely used to study HBx, exhibit pronounced aneuploidy, chromosomal instability, and altered proteomes compared to primary hepatocytes (23, 24). In addition, they already display tumour-derived transcriptional programs, limiting their application to study HBx-associated tumorigenesis. While primary human hepatocytes (PHH) provide a more physiologically relevant system, their use is constrained by rapid dedifferentiation, limited lifespan, donor-to-donor variability, and limited amenability to genetic manipulation (25, 26). Similarly, induced pluripotent stem cell-derived hepatocyte-like cells retain foetal-like transcriptional signatures and lack full functional maturation (27, 28). Conversely, *in vivo* animal models, while providing systemic insights, fail to recapitulate human-specific physiological and pathological responses (29).

Adult human liver-derived organoids provide a promising alternative, enabling long-term expansion of genetically stable, three-dimensional (3D) cultures derived from human adult stem cells. These organoids recapitulate key features of native liver tissue, including cellular heterogeneity, metabolic function, and physiologically relevant stress responses. However, efficient differentiation of these bipotent progenitor cells into fully mature hepatocyte-like cells remains a major challenge, highlighting the need for improved culture conditions (30).

In this study, we leverage liver organoids derived from adult human livers to investigate HBx function in hepatocytes. We first implement and characterize a differentiation medium that promotes the generation of more mature hepatocyte-like cells. Using this system, we systematically address key variables influencing HBx activity, including expression level, donor background, and protein tagging. To this end, we employ multiple donor-derived organoid lines and complementary expression strategies, enabling both cognate and inducible HBx expression fused to distinct protein tags. Using this approach, we demonstrate that HBx reshapes hepatocyte identity and apoptotic signalling to promote hepatocyte survival. Together, our organoid-based platform provides a relevant system to elucidate HBx-mediated mechanisms underlying HBV persistence and liver tumorigenesis.

## Results

### DM+ condition promotes the generation of hepatocyte-like organoids with enhanced differentiation

Adult stem cell (ASC)-derived liver organoids are enriched for bipotent liver progenitor-like cells under expansion conditions (expansion medium; EM), supporting self-renewal and scalability. Upon differentiation (differentiation medium; DM), we have previously demonstrated that these hepatocyte-like organoids (HLOs) are susceptible to HBV infection and support viral replication (31, 32). However, DM-HLOs exhibit only partial hepatocyte maturation, leaving key metabolic, structural, and functional properties required to recapitulate adult liver physiology underdeveloped (30). Therefore, we implemented a modified differentiation medium based on DM, termed enhanced differentiation medium (DM+), which was reported to improve terminal differentiation of bipotent progenitor cells present in EM-HLOs toward hepatocytes (33). Given the central role of Wnt signalling in liver development, regeneration, zonation, and homeostasis (34, 35), a number of small molecules modulating this pathway are added to this formulation. In addition, motivated by emerging evidence implicating acetylcholine signalling in hepatocyte proliferation, metabolic regulation, and liver pathology (36, 37), carbachol, a stable acetylcholine analogue is also included. Next, we compared DM+ to DM for its ability to promote hepatocyte-like differentiation, and characterized the degree of terminal differentiation achieved in our system (Fig. 1A-D).

**Fig. 1.**
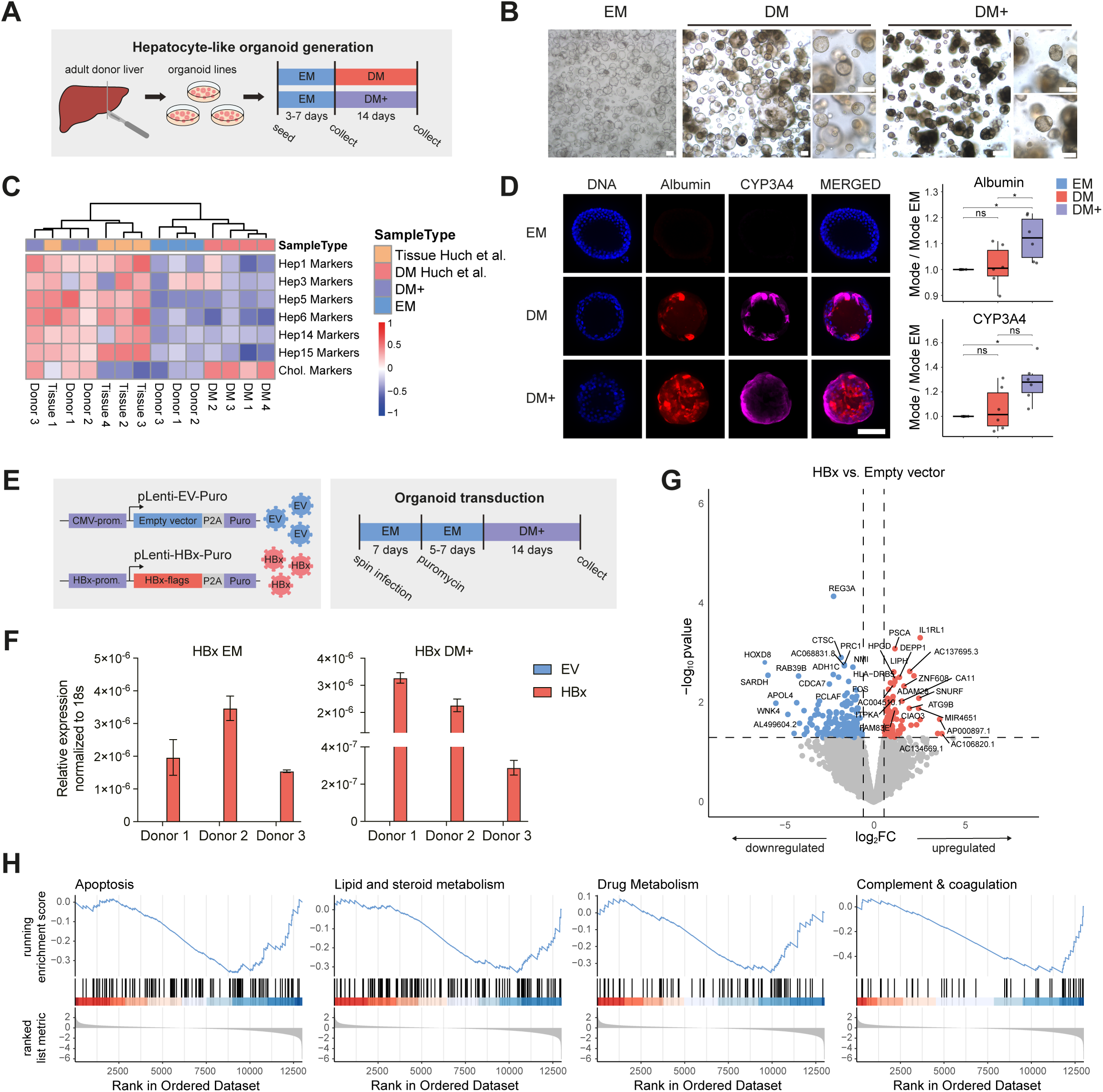
DM+ condition promotes the generation of hepatocyte-like organoids with enhanced differentiation and enables cognate promoter driven HBx expression in hepatocyte-like organoids. (A) Schematic of HLO generation: adult human liver-derived organoids were expanded in EM and differentiated for 14 days in DM/DM+. (B) Representative brightfield images before and after differentiation (donor 8; scale bar, 200 μm). (C) GSVA of hepatocyte/cholangiocyte marker sets (38)using bulk RNA-seq. and published microarray data (30). Samples were hierarchically clustered based on enrichment profiles. (D) Immunofluorescence before/after differentiation (maximum projection, donor 11; scale bar, 100 μm) and quantification of mean fluorescence intensity per donor; each dot represents the mode of the per-organoid mean intensities for an individual donor, normalized to the EM condition. Statistical significance was assessed using a paired Wilcoxon signed-rank test. (E) Lentiviral HBx construct and experimental design. (F) HBx mRNA expression by RT-qPCR. (G) Differential gene expression between empty vector and HBx-expressing DM+-HLOs. Significant DEGs were defined by p value < 0.05 and |log₂ fold change| > 0.58. (H) GSEA showing negative enrichment of apoptosis and liver-function gene sets in HBx-expressing DM+-HLOs.

Following a 14-day differentiation, organoids displayed marked morphological changes including increased darkening and compaction, which were most prominent in DM+ (Fig. 1B). Both DM and DM+ induced expression of mature hepatocyte markers (Fig. S1A). However, organoids cultured in DM+ showed a more pronounced reduction in the progenitor and ductal marker SOX9. Notably, while differentiation in DM triggered upregulation of the foetal hepatocyte marker AFP, organoids differentiated in DM+ did not exhibit this induction (Fig. S1A). To obtain a broader view of transcriptional identity and to enable comparison with primary liver tissue, we performed gene set variation analysis (GSVA) using hepatocyte and cholangiocyte gene sets (38). Enrichment was assessed using bulk RNA-sequencing (RNA-seq.) data from organoids in EM and DM+, integrated with microarray datasets from primary human liver tissue and organoids differentiated in DM (30). As expected, organoids in EM showed minimal enrichment for hepatocyte-associated signatures. By contrast, organoids differentiated in DM+ exhibited a clear increase in hepatocyte-like gene enrichment, clustering closely with primary liver samples. Organoids differentiated in DM, however, did not display this hepatocyte-like enrichment (Fig. 1C). Consistent with these findings, DM+ also yielded the strongest induction of mature hepatocyte genes at the protein level (Fig. 1D), confirming improved generation of hepatocyte-like organoids in the novel medium formulation.

Previously, it has been reported that exposure of undifferentiated organoids to BMP7, prior to differentiation, improves differentiation efficiency (30). Implementation of this step prior to differentiation in DM+, however, did not result in improved differentiation (Fig. S1B-C) and was excluded in subsequent experiments. DM+-HLOs consistently showed robust expression of mature hepatocyte markers and downregulation of ductal/progenitor markers at the protein level (Fig. 1D, Fig. S1B-1C). Functionally, DM+-HLOs demonstrated increased glycogen storage capacity (Fig. S1D). Differential gene expression analysis between EM and DM+ revealed upregulation of hepatocyte-associated metabolic pathways and downregulation of cell-cycle and stem-cell related signalling programs (Fig. S1E-G). Taken together, these findings indicate that DM+ promotes the consistent generation of hepatocyte-like organoids with enhanced differentiation.

### Exogenous expression of HBx in DM+-HLOs results in anti-apoptotic and dedifferentiated transcriptomic signatures

To dissect the mechanistic consequences of HBx expression and interrogate HBx-dependent pathways and cellular responses in an adult hepatocyte context, we generated transgenically modified normal HLOs exogenously expressing HBx. First, we generated a lentiviral construct containing the 2x flag tagged HBx coding sequence under the control of the cognate HBx promoter followed by a puromycin selection marker (pLenti-HBx-Puro) (Fig. 1E). Expression of lentivirally delivered HBx and the empty vector control (pLenti-EV-Puro) was validated in HepG2 cells at both the RNA and protein levels (Fig. S2A-B), and used to generate stably transduced HLOs mimicking the low HBx expression levels in HBV infected hepatocytes, controlled by the HBx cognate promoter. Stably transduced HLOs were generated and expanded in EM, and subsequently differentiated into DM+-HLOs (Fig. 1E). Total RNA was isolated from transgenic organoids maintained in either EM or DM+ to quantify HBx expression levels (Fig. 1F), and samples were subjected to 3′ mRNA sequencing to assess global transcriptomic changes consequent to HBx expression in DM+-HLOs. RNA-seq. analysis identified 293 differentially expressed genes (DEGs), of which 120 were upregulated and 173 were downregulated in HBx-expressing HLOs compared to controls (Fig. 1G). To explore HBx function in our hepatocyte-like organoid system, we performed gene set enrichment analysis (GSEA) using KEGG pathways. HBx expression in DM+-HLOs resulted in negative enrichment of apoptosis-associated gene sets (Fig. 1H). In addition, critical hepatocyte functions, such as metabolism, detoxification, and the synthesis of complement and coagulation factors (39–42), showed negative enrichment in HBx-expressing organoids (Fig. 1H), suggesting impaired acquisition or maintenance of hepatocyte identity.

We next sought to validate our observations in primary human liver samples. To this end, we analysed a bulk RNA-seq. dataset derived from liver samples in the TCGA Research Network (https://www.cancer.gov/tcga). First, reads were aligned to a combined human-HBV reference genome to detect viral transcripts, and samples were classified as HBV-infected or non-infected based on total HBx (X) and S gene expression. Differential gene expression followed by GSEA confirmed downregulation of apoptosis and liver-function gene sets in HBV-infected samples (Fig. S2C-D), consistent with our findings in DM+-HLOs. As expected, HBx and S expression showed high positive correlation (Fig. S2E). Additionally, given the presence of integration-competent HBV and an immune compartment in primary samples, HBx expression was associated with increased *TERT* expression (Fig. S2F) and negatively correlated with immune-associated genes (Fig. S2E), suggesting HBV integration and suppression of antigen presentation, respectively.

Although the cognate promoter-driven model enabled HBx expression in hepatocyte-like organoids at physiologically relevant levels, it relied on generation of constitutive HBx expression in organoids under expansion conditions which remained expressed during differentiation towards DM+-HLOs. To determine whether HBx reduces hepatocyte identity by interfering with the differentiation process itself or by inducing dedifferentiation after the differentiated hepatocyte state is established, and to better characterize the direct mechanistic effects of HBx, we next developed a Tet-On inducible system enabling temporal control of HBx expression and its induction specifically in DM+-HLOs. To inducibly express HBx, two lentiviral constructs were generated, tagging HBx with either a V5 epitope or EGFP (pLenti-HBx-V5-Puro and pLenti-HBx-EGFP-Puro). The distinct tags enabled both immunofluorescence-based detection and live tracking of HBx expression using the V5 and EGFP variant, respectively. In addition, use of constructs with distinct tags allowed identification and assessment of common HBx-induced mechanisms present independently of the tag, which could potentially interfere with HBx interactions.

Lentiviral particles were then generated and validated in HepG2 cells (Fig. S3A). To verify that HBx induction was strictly transgene-dependent, wild-type HepG2 cells were treated with doxycycline (dox.), which did not result in detectable HBx expression (Fig. S3B, S3D), while dox. treatment in transduced HepG2 cells induced HBx expression in a dose- and time-dependent manner, with induction plateauing at approximately 2 µg/mL (Fig. S3C-D). This concentration was therefore used for all subsequent organoid experiments. Next, EM-HLOs were transduced with either the EGFP- or V5-tagged inducible construct, followed by puromycin selection to obtain stable lines, which were then differentiated toward DM+-HLO (Fig. 2A). HBx expression was induced by addition of 2 µg/mL dox. for 24 hours (h), and induction was confirmed at both the RNA and protein levels (Fig. 2B, Fig. S3E). Once again, it was verified that HBx induction is strictly transgene dependent and inducible, with no detectable HBx expression from parental (non-transduced) organoid lines in the presence of dox. (Fig. S3F). Compared to the expression levels obtained by the cognate HBx promoter driven system, the Tet-On inducible system resulted in markedly higher levels of HBx expression in both EM and DM+ (Fig. S3G).

**Figure 2.**
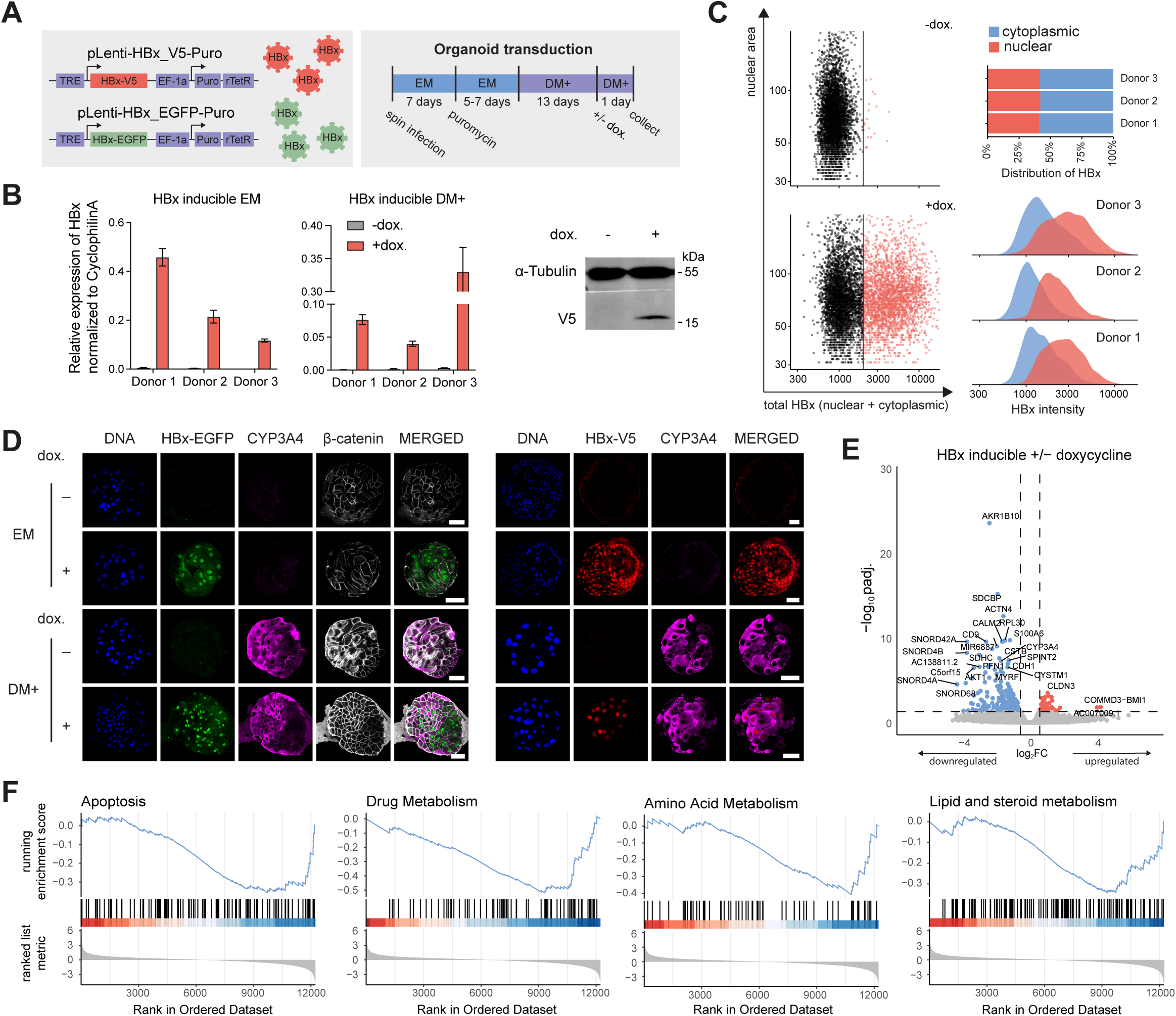
Exogenous expression of HBx in DM+-HLOs results in anti-apoptotic and dedifferentiation transcriptomic signatures. (A) Tet-on lentiviral HBx constructs (V5- or EGFP-tagged) and experimental design; HLOs were transduced, selected, differentiated for 14 days, and induced with dox. for 24 h. (B) Left, HBx mRNA expression and, right, Western blot validation. (C) Left, representative scatterplot of total HBx intensity against nuclear area of cells in DM+-HLOs (donor 1). Vertical red line indicates the HBx intensity threshold used to detect HBx-positive cells. Top right, total HBx abundance detected in the nuclear and cytoplasmic compartment of the cell, determined using integrated intensity. Bottom right, overlaid density plots comparing the cytoplasmic and nuclear HBx intensity. (D) Representative immunofluorescence images before/after differentiation ± dox (donor 2; scale bar, 50 µm). Merged channel is composed of all channels except DNA. (E) Differential gene expression between DM+-HLOs ± dox. Significant DEGs were defined by adjusted p value < 0.05 and |log₂ fold change| > 0.58. (F) GSEA showing negative enrichment of apoptosis and liver-function gene sets in HBx-expressing organoids.

Similar to its reported functions, the subcellular localization of HBx has been reported to vary depending on cellular model and expression levels. To assess HBx localization in DM+-HLOs, we performed fluorescence microscopy on pLenti-HBx-EGFP-Puro transduced HBx-expressing DM+-HLOs after 24 h dox. treatment. Nuclear and cytoplasmic compartments were segmented for each cell, and cells in dox.-treated samples were classified as HBx-positive or HBx-negative based on mean total HBx intensity (Fig. 2C). In HBx-positive cells, both nuclear and cytoplasmic HBx fractions were quantified. Although total HBx abundance, as measured by integrated intensity, was greater in the cytoplasm across donors, nuclear HBx showed higher concentration, reflected by stronger nuclear signal intensity (Fig. 2C). This localization was consistent across differentiated and undifferentiated organoids, independent of the tag used to visualize HBx (Fig. 2D).

It has previously been reported that nuclear HBx acts as a co-activator and modifies the host epigenome through regulating the interaction of chromatin modifying enzymes and transcription factors with host chromatin (16–18). Additionally, HBx is thought to activate several signalling pathways in the cytoplasm to modulate host gene transcription (43). To investigate the impact of HBx on chromatin accessibility we performed assay of transposase-accessible chromatin with sequencing (ATAC-seq.) (44) on differentiated, transduced DM+-HLOs treated with or without dox. to express HBx. We found that 610 chromatin regions were differentially accessible (DARs) between HBx-expressing and non-expressing DM+-HLOs of which the majority (68%) displayed reduced accessibility upon HBx expression (Fig. S4A). We used the Genomic Regions Enrichment of Annotations Tool (GREAT) (45) to assign potential genes regulated by these DARs and found that the majority of these DARs are located distally (>5kb) from transcription start sites (Fig. S4B). Motif analysis revealed a significant enrichment of binding motifs for the AP1-family, FOX-family and HNF4α transcription factors in the DARs (Fig. S4C). Analysis of ChIP-seq data generated by the ENCODE consortium (46) revealed substantial (>20%) overlap of the DARs with binding regions of FOSL2, FOXA2, FOXA3 and FOXO1 in HepG2 cells while overlap with HNF4α was found in liver samples (Fig. S4D).

We next performed RNA sequencing on differentiated, transduced DM+-HLOs treated with or without dox. to express HBx (Fig. 2A). To account for doxycycline-induced off-target effects, parental organoid lines were also profiled with and without dox. treatment. Differential gene expression analysis revealed that dox. alone resulted in only a limited number of transcriptomic changes (3 upregulated DEGs) (Fig. S3H). Following HBx expression in transduced DM+-HLOs, 243 DEGs were identified, 66 of which were upregulated and 177 were downregulated compared to controls (Fig. 2E). When we compared the DEGs with the genes assigned to the DARs by GREAT we found only limited overlap between the gene sets (Fig. S4E), likely reflecting that chromatin accessibility changes at distal regulatory elements may precede or occur independently of transcriptional changes in associated genes. Consistent with our results using the cognate promoter-driven HBx construct and with observations in infected primary patient samples, dox. inducible HBx expression negatively enriched apoptosis- and liver function-related pathways cross 3 independent donor liver organoid lines (Fig. 2F). Additionally, KEGG pathway enrichment on DEGs in response to HBx expression further supported downregulation of apoptosis, and hallmark liver metabolic pathways (Fig. S3I). Moreover, genes involved in protein processing in the endoplasmic reticulum (ER) and the proteasome were downregulated (Fig. S3I).

Because HBx consistently downregulated apoptosis- and liver function-associated genes across primary patient samples, different organoid lines, tagging strategies, expression levels, and induction durations, we next examined how these transcriptomic alterations translate into functional changes in hepatocyte-like organoid behaviour. The Tet-On inducible system was used in all subsequent functional experiments, as it enables temporal control of HBx expression, allowing assessment of direct transcriptional targets (24 h post-induction), and restricts induction to differentiated DM+-HLOs, providing a more physiologically relevant context for modelling HBx function during HBV infection.

### Exogenous expression of HBx in DM+-HLOs attenuates the induction of apoptosis

In addition to gene set-wide alterations in apoptosis-related pathways, expression of individual genes across multiple apoptotic programs was significantly reduced by exogenous expression of HBx across donor lines (Fig. 3A). Notably, caspase 9 showed substantial downregulation in response to HBx expression (Fig. 3A). Caspase 9 is an initiator of the intrinsic- or mitochondrial apoptosis pathway, whose activation leads to downstream cleavage and activation of caspase 3 and 7, which are responsible for initiation of further hallmarks of apoptosis (47, 48). To functionally assess how HBx expression affects apoptosis, we used the dox.-inducible Tet-On system in transduced organoids to express HBx for 24h in DM+-HLOs. Subsequently, we induced apoptosis in the DM+-HLOs via a caspase 9 dependent mechanism by treatment with α-(trichloromethyl)-4-pyridineethanol (PETCM) for 12h, a small molecule that promotes apoptosome formation, leading to caspase 9 activation (49) (Fig. 3B). We examined the effect of HBx expression on downstream caspase 3/7 activation by subjecting the samples to high-content imaging, visualizing and quantifying activated caspase 3/7, HBx, and DNA per individual nucleus (Fig. S5A). Assay specificity and analysis performance were validated by confirming robust induction of HBx in response to dox., as well as caspase 3/7 activation in response to treatment with increasing concentrations of the caspase 9 inducer PETCM across all donors (Fig. 3C). At the single-cell level, PETCM treatment resulted in the appearance of a distinct caspase 3/7-high population (y-axis, Fig. 3D). Importantly, the cells exhibiting the strongest caspase 3/7 activation were predominantly the HBx-low subpopulation, while cells expressing high levels of HBx showed low caspase 3/7 activation (Fig. 3D), consistent with the notion that HBx expression attenuates apoptosis.

**Figure 3.**
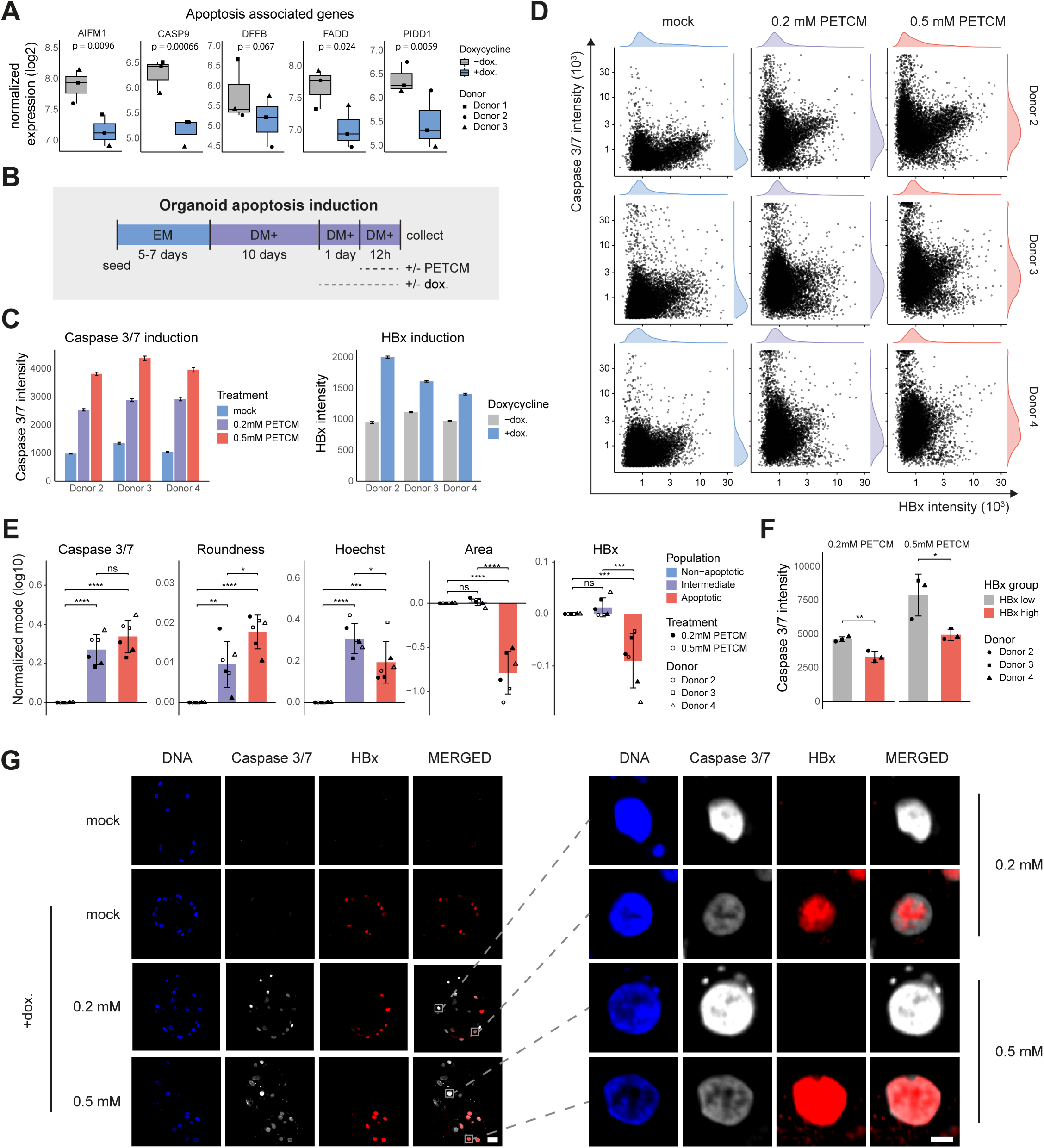
Exogenous expression of HBx in DM+-HLOs attenuates the induction of apoptosis. (A) RNA-seq. analysis of apoptosis-associated genes in DM+-HLOs ± dox. P-values derived from a Wald test in DESeq2. (B) Experimental design: HLOs differentiated for 10 days, induced ± dox for 24 h, then exposed to 0, 0.2, or 0.5 mM PETCM for 12 h. (C) Mean activated caspase 3/7 intensity per cell ± SEM per treatment condition per donor; n > 8000 individual cells per condition. (D) Scatterplots of single-cell HBx versus caspase 3/7 intensity obtained from DM+-HLOs +dox. (E) Mean of normalized modes ± SD. Data is normalized, per donor-treatment combination, to the non-apoptotic population. Statistical significance was assessed using one-way ANOVA followed by Tukey’s multiple comparisons test. (F) Caspase 3/7 intensity in apoptotic HBx-high versus HBx-low cells. Statistical significance was assessed using one-way ANOVA followed by Tukey’s multiple comparisons test. (G) Representative immunofluorescence images ± dox with PETCM (donor 2; scale bar, 25 (left) and 5 µm (right)). Merged channel is composed of all channels except DNA.

To better understand the relationship between HBx expression and induction of apoptosis, the total cell population of each donor DM+-HLO-treatment condition was stratified into non-apoptotic, intermediate, and apoptotic groups. These groups were defined using canonical apoptotic features including caspase 3/7 activity and cell size (Fig. S5B) (50–52). Intermediate and apoptotic cells displayed increased caspase 3/7 activation and increased roundness relative to non-apoptotic cells (Fig. 3E, Fig. S5C), while also showing elevated Hoechst intensity, consistent with apoptotic chromatin condensation (Fig. 3E, Fig. S5C) (50–52). Apoptotic cells further exhibited reduced nuclear area, aligning with the nuclear shrinkage and fragmentation characteristic of apoptosis (Fig. 3E, Fig. S5C) (50–52). Importantly, apoptotic nuclei showed markedly reduced HBx abundance compared to non-apoptotic cells (Fig. 3E, Fig. S5C).

After classifying apoptotic states across all samples, each apoptotic population was further divided into HBx-high and HBx-low subpopulations, based on whether HBx expression levels fell within the upper or lower half of the distribution for that donor-treatment condition (Fig. S5D). Within the apoptotic population, HBx expression once again correlated with reduced caspase 3/7 activation (Fig. 3F-G). Taken together, these findings directly link HBx expression to decreased induction of apoptosis in hepatocyte-like organoids.

### Expression of HBx results in dedifferentiation of hepatocyte-like organoids and evasion of apoptosis

Having established that HBx attenuates the induction of apoptosis, we next examined how HBx affects hepatocyte identity. Similar to its effect on apoptosis-related genes, across donor lines HBx expression also negatively regulated the expression of individual liver function-associated genes, in addition to the broader gene set-wide reduction captured by GSEA (Fig. 2F, Fig. 4A). Notably, CYP3A4, the most abundant drug-metabolizing cytochrome P450 enzyme in the human liver and a hallmark of hepatocyte maturation, was significantly downregulated in HBx-expressing organoids (Fig. 4A) (53).

**Figure 4.**
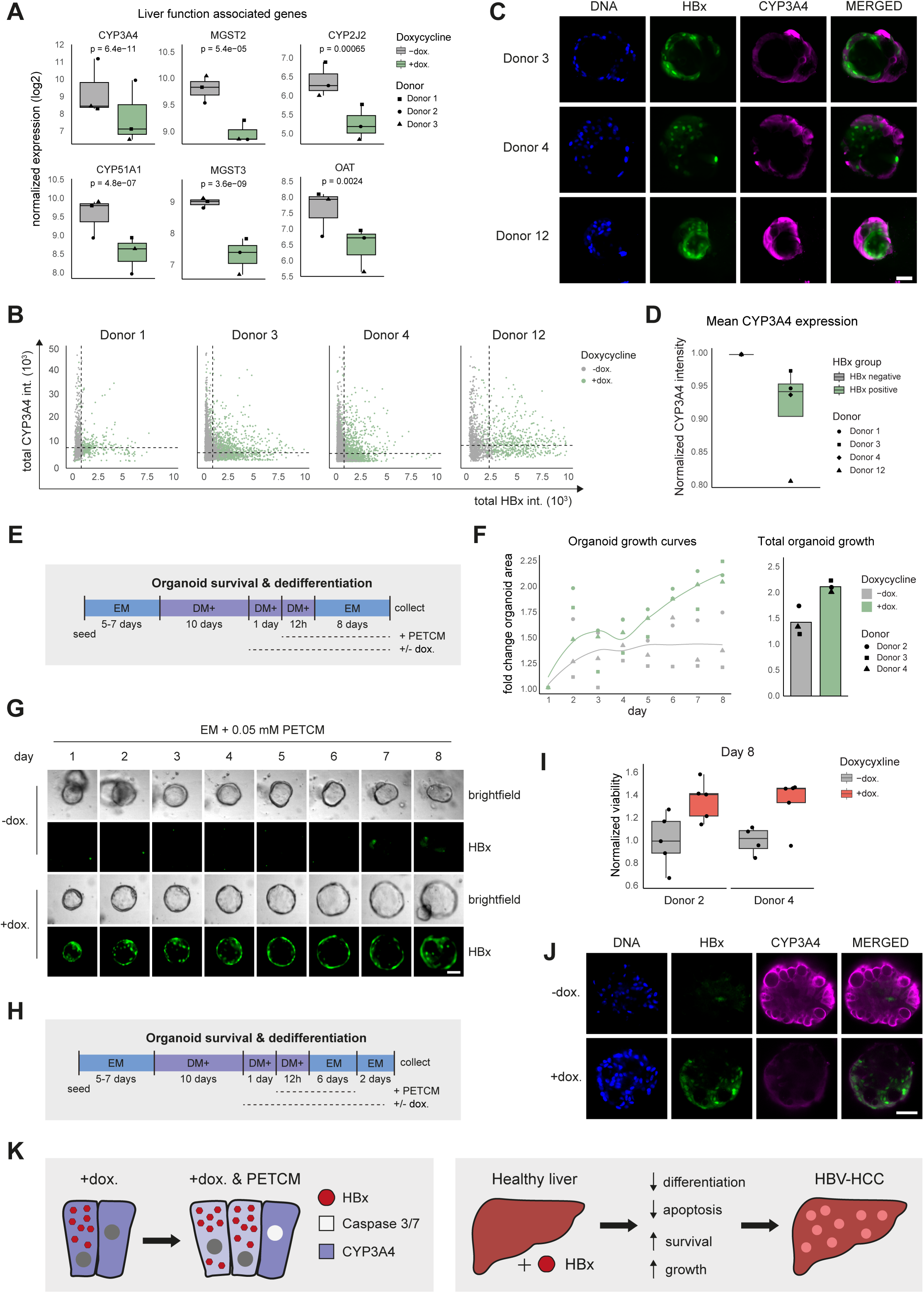
Exogenous expression of HBx results in dedifferentiation of DM+-HLOs and evasion of apoptosis. (A) RNA-seq. analysis of liver function-associated genes in DM+-HLOs ± dox. P-values derived from a Wald test in DESeq2. (B) Scatterplots of single-cell HBx versus CYP3A4 intensity. Vertical and horizontal dotted lines indicate the gating used to determine HBx- and CYP3A4-positivity, respectively. (C) Representative immunofluorescence of +dox. organoids (scale bar, 50 µm). Merged channel is composed of all channels except DNA. (D) Normalized mean CYP3A4 intensity in HBx-negative versus HBx-positive cells in +dox. DM+-HLOs. (E) Experimental design corresponding to Fig. 4F-G. (F) Organoid growth during PETCM [0.05 mM] treatment in EM; n ≥ 3 organoids per biological replicate, represented as the mean across donors. (G) Representative images of organoids during treatment (donor 4; scale bar, 50 µm). (H) Experimental design corresponding to Fig. 4I-J and S6E-F. (I) Normalized viability at day 8 after re-seeding in EM. Viability was normalized to the -dox. condition and determined using Cell-titer Glo 3D. (J) Representative immunofluorescence staining of organoids at day 8 after re-seeding in EM. Merged channel is composed of all channels except DNA (donor 2; scale bar, 50 µm). (K) Schematic representation of the proposed oncogenic mechanisms of HBx. Exogenous HBx expression in hepatocyte-like cells induces dedifferentiation, and suppresses apoptosis, conferring a proliferative advantage under challenge conditions, thereby potentially contributing to the development of HBV-associated HCC.

To investigate the relationship between HBx expression and hepatocyte differentiation at the single-cell level, HBx transgenic DM+-HLOs with or without doxycycline were processed as previously described (Fig. 2A), and imaged using high-content microscopy in which DNA, HBx and CYP3A4 were visualized. HBx and CYP3A4 intensities were quantified within both the nucleus and the whole-cell outline (Fig. S6A). Before analysis, debris and mis-segmented objects were excluded based on nuclear size, shape, and Hoechst signal (Fig. S6C).

Scatterplots of total HBx versus total CYP3A4 intensity revealed robust HBx induction upon dox. treatment, as expected (x-axis, Fig. 4B). Intriguingly, CYP3A4-high cells predominantly arose from the HBx-low fraction, suggesting an inverse relationship between HBx expression levels and hepatocyte differentiation (Fig. 4B). To quantify this, mature hepatocyte-like cells were defined using donor-matched thresholds derived from EM-HLOs. For each donor, the 90th percentile of CYP3A4 expression in EM-HLO cells was used as a cutoff, and cells in DM+-HLOs exceeding this threshold were classified as CYP3A4-positive (Fig. S6B), representing approximately 45% of the population. This 45% fraction was subsequently used to define the mature hepatocyte compartment in DM+-HLOs. Within this mature population, HBx-positive and HBx-negative cells were classified using the basal minus dox. condition as a reference (Fig. 4B). Across all donors, HBx-high mature hepatocyte-like cells consistently exhibited reduced CYP3A4 expression (Fig. 4C-D), in agreement with a role for HBx in hepatocyte de-differentiation.

As noted earlier, DM+-HLOs show reduced proliferation and diminished stem cell-associated signatures compared to EM-HLOs (Fig. S1E-G). This shift is accompanied by a halt in proliferation (Fig. S6D). *In vivo*, the adult human liver can regenerate following resection, in part through normally quiescent hepatocytes re-entering the cell cycle (54, 55). Mature hepatocytes also possess the capacity to dedifferentiate toward progenitor-like states under specific conditions (56, 57). We therefore asked whether re-exposing HBx transgenic DM+-HLOs to expansion conditions could trigger dedifferentiation, and whether HBx expression might promote this process while simultaneously reducing apoptosis.

To test this, DM+-HLOs were treated with or without dox. for 24h to express HBx before exposure to the caspase 9 inducer PETCM. After 12 hours of PETCM treatment, organoids were transferred into EM to promote proliferation, in the presence of PETCM and dox. maintained throughout the culture period to ensure induction of apoptosis concomitant with continuous expression of HBx (Fig. 4E).

Upon transfer to EM, HBx transgenic DM+-HLOs regained proliferative capacity, reflected by an increase in total organoid area detected in brightfield images (Fig. 4F-G). In organoids lacking HBx induction, this expansion plateaued (Fig. 4F-G), likely due to the pro-apoptotic effects of PETCM. Remarkably, HBx-expressing organoids not only showed enhanced dedifferentiation capacity, as indicated by greater increases in total organoid area, but also sustained proliferation for an extended period in the continued presence of PETCM (Fig. 4F-G). These effects were consistent with increased overall viability and lower levels of CYP3A4 expression compared to organoids not expressing HBx (Fig. 4H-J, Fig. S6E-F). Thus, HBx expression supports cell survival by both promoting hepatocyte de-differentiation and by attenuating apoptosis, which suggests that HBx may similarly suppress hepatocytes differentiation and apoptosis while enhancing survival and proliferation during HBV infection, thereby contributing to the pathogenesis of HBV-associated HCC (Fig. 4K).

## Discussion

Viruses have evolved diverse strategies to exploit their host’s cellular machinery for their own propagation, many of which overlap with canonical hallmarks of cancer. Processes such as immune evasion, suppression of apoptosis, and deregulation of cell cycle progression confer clear advantages in both viral infection and tumour evolution (58–60). In the context of HBV, these parallels are particularly evident. Integration of the HBV genome into host DNA contributes to genomic instability, alters host gene expression, and increases mutational burden (8). Consistent with previous reports of recurrent HBV integration near the *TERT* locus, we observed increasing *TERT* expression with higher levels of HBx expression in our patient cohort (Fig. S2F), supporting a mechanism for replicative immortality shared with many cancers (61, 62). In addition to these integration-driven effects, viral proteins themselves can directly modulate host cell behaviour. For instance, higher HBx expression in our patient cohort was associated with reduced expression of immune-related genes, suggesting that HBx may contribute to an immunosuppressive environment (Fig. S2E). Better understanding of the oncogenic functions and underlying mechanisms of the viral proteins is pivotal for developing targeted interventions against HBV-associated HCC. Here, we specifically investigated the oncogenic properties of the HBV-encoded HBx protein and identified loss of hepatocyte identity and repression of apoptosis as key cellular responses to its expression.

A major challenge in defining HBx function is the variability of its reported effects, including on apoptosis, which is largely attributable to differences in experimental systems, expression levels, and cellular backgrounds (63–65). In hepatoma cell lines such as HepG2 and Huh7, HBx has been reported to promote apoptosis through mechanisms including c-FLIP sequestration and TRAIL receptor signalling (66–68).

However, anti-apoptotic effects have also been described in similar and related cell systems, including via inhibition of FAS-mediated apoptosis and activation of PI3K/AKT signalling (69–71). In primary human and rodent hepatocytes, HBx has been shown to inhibit apoptosis via FAS signalling and promote survival through AKT or Wnt/β-catenin signalling (72–75). Thus, opposing pro- and anti-apoptotic functions of HBx have been reported indicating that the role of HBx in apoptosis is highly context-dependent.

In addition to its effects on apoptosis, HBx has been reported to modulate hepatocyte hallmark processes such as glucose, lipid, and amino acid metabolism, and has been implicated in regulating cellular plasticity (76–78). However, its impact on hepatocyte identity remains under debate. On the one hand, HBx has been shown to suppress differentiation-associated programs, for example through downregulation of HNF4α, leading to reduced expression of targets such as CYP2E1 (79, 80). Given that HNF4α is a key regulator of hepatocyte identity and controls genes including CYP3A4, these findings support a model in which HBx impairs hepatocyte differentiation (81). On the other hand, HBx has also been reported to increase CYP3A4 expression via activation of PXR signalling in hepatoma cell lines and *in vivo* models, suggesting context-dependent or even opposing effects on hepatocyte-specific functions (82, 83). Thus, similar to its role in apoptosis, the impact of HBx on hepatocyte identity appears highly variable across experimental systems, underscoring the need for physiologically relevant models to resolve its effects in differentiated human hepatocytes.

To address these limitations, we employed a human liver organoid model derived from adult stem cells and we characterized improved differentiation conditions to generate more mature hepatocyte-like cells. This system preserves key aspects of hepatocyte biology while allowing controlled manipulation of HBx expression across multiple donor backgrounds. Using this platform, we demonstrate that HBx promotes a coordinated shift in hepatocyte state characterized by increased resistance to apoptosis and reduced lineage fidelity. While the role of HBx in apoptosis has been debated, our findings in a differentiated hepatocyte context support a predominantly anti-apoptotic function. Interestingly, it has previously been reported that HBx inhibits apoptosis via modulation of the ER stress response (84), which was also found to be dysregulated in our analysis (Fig. S3I). In parallel, HBx expression led to a reduction in hepatocyte-specific features. Importantly, these effects occurred concurrently, suggesting that HBx uncouples hepatocyte survival from differentiation. Such a state may allow damaged or dysregulated hepatocytes to evade elimination while acquiring a more permissive, proliferative phenotype, thereby contributing to early stages of HBV-associated tumorigenesis.

Mechanistically, HBx is not thought to exert its effects through direct binding to DNA, but rather through interactions with host proteins that regulate transcription, signalling, and chromatin dynamics (16–20, 80, 85). It has been proposed that the subcellular compartmentalization of HBx drives its distinct functions, with cytoplasmically localized HBx engaging signalling cascades, while nuclear HBx modulating chromatin associated complexes and transcription factor activity, dysregulating host gene expression (86). HBx localization has also been shown to depend on its expression level, with lower expression favouring nuclear localization and higher expression driving its accumulation in the cytoplasm (86–89). In line with this, our Tet-On inducible system, which achieved higher HBx expression levels than the system driven by the cognate HBx promoter, showed a predominantly cytoplasmic HBx localization, although signal intensity was markedly enriched in the nucleus. This nuclear localization suggests that its effects on hepatocyte identity and survival may be mediated through modulation of transcriptional programs via interaction with transcription regulatory complexes. This interaction-driven mode of action provides a plausible explanation for the context-dependent effects of HBx observed across different systems. In this framework, the coordinated suppression of apoptosis and loss of hepatocyte identity observed in our model may arise from HBx-mediated rewiring of chromatin-associated regulatory networks rather than direct transcriptional control.

Our ATAC-seq. data support this model by showing that HBx expression is associated with widespread changes in chromatin accessibility, predominantly at distal regulatory regions, consistent with indirect modulation of host transcriptional programs rather than direct DNA binding by HBx. The enrichment of motifs for HNF4α, FOX family members, and AP-1 factors suggests that HBx may alter chromatin states at regulatory elements controlled by transcription factors that are central to both hepatocyte identity and cellular stress responses. In particular, HNF4α is a key regulator of hepatocyte differentiation, and disruption of its regulatory network provides a plausible mechanism for the loss of hepatocyte-specific features observed in our model. The enrichment of AP-1 motifs is also notable given its role in hepatocyte priming during liver regeneration, where it promotes expression of proliferative programs while repressing liver-specific differentiation genes, and its involvement in the regulation of apoptosis (90, 91). Future studies combining chromatin accessibility profiling with mapping of HBx-associated protein complexes in differentiated human hepatocytes will be important to further resolve how HBx reshapes the epigenetic and transcriptional landscape.

Although our organoid system overcomes several substantial limitations of traditional models, providing a strong platform to investigate hepatocyte-intrinsic effects of HBx, it does not recapitulate the influence of non-epithelial liver cell populations. In particular, the absence of non-parenchymal cell types, including immune cells, hepatic stellate cells, endothelial cells, and Kupffer cells, limits the ability to capture intercellular signalling processes that contribute to chronic inflammation, fibrogenesis, and immune modulation during disease progression. Immune evasion is a key hallmark of cancer and has been observed in HBV-associated disease, for example through modulation of MHC class I expression (58, 59, 92), and HBx itself has also been implicated in promoting such immune evasion phenotypes (93, 94). While hepatocytes themselves express MHC-I, its regulation is highly dependent on cytokine signalling, which is largely mediated by non-parenchymal cells such as Kupffer cells and other immune populations (95–97). In addition to immune evasion, these broader microenvironmental cues are known to shape hepatocyte plasticity, survival, and malignant transformation, their absence may therefore limit the extent to which HBx-driven disease-relevant phenotypes can be fully modelled in our system. Future co-culture organoid systems incorporating these components will therefore be important to further define how HBx contributes to hepatocellular carcinoma development *in vivo*.

In conclusion, our findings support a model in which HBx acts as a central modulator of hepatocyte state by uncoupling cellular survival from lineage fidelity, thereby contributing to the pathogenesis of HBV-associated HCC. Using a physiologically relevant human liver organoid system, we show that HBx-driven repression of apoptosis and loss of hepatocyte identity are consistently observed across donors, tagging strategies, and expression levels (Fig. 4K). These transcriptomic changes are accompanied by widespread remodelling of chromatin accessibility at distal regulatory elements, enriched for key transcription factor networks governing hepatocyte differentiation and stress responses. Future work defining the chromatin-associated protein complexes interacting with HBx will be essential to elucidate how it indirectly reprograms host epigenetic and transcriptional states. While these effects are likely further shaped by the hepatic microenvironment *in vivo*, our study defines a hepatocyte-intrinsic framework for HBx function and provides mechanistic insight into how viral perturbation of epithelial state may contribute to liver tumorigenesis.

## Abbreviations

HBx: Hepatitis B virus protein X
HBV: Hepatitis B virus
HCC: hepatocellular carcinoma
PHH: primary human hepatocytes
HLOs: hepatocyte-like organoids
ASC: adult stem cell
EM: expansion medium
DM: differentiated medium
DM+: enhanced differentiation medium
RNA-seq: RNA sequencing
GSEA: gene set enrichment analysis
TCGA: The Cancer Genome Atlas
dox: doxycycline
PCR: Polymerase Chain Reaction
RT-qPCR: Reverse Transcription-quantitative PCR
DEG: differentially expressed gene
DAR: differentially accessible region
ATAC-seq: assay of transposase-accessible chromatin with sequencing

## Acknowledgements

We gratefully acknowledge Adam Karpf, Alice Ting & Didier Trono for providing the TLCV2, dCas13d-dsRBD-APEX2, pMD2.G and psPAX2 plasmids. The Fleming Genomics Facility is supported by project SingleOut (HFRI-FM17C3-3780) funded by the Hellenic Foundation for Research and Innovation (H.F.R.I.), project MIS 6004752 funded by the Regional Operational Programme ‘ATTICA’ (NSRF 2021–2027) and co-financed by Greece and the European Union (European Regional Development Fund), and the National Recovery and Resilience Plan Greece 2.0, funded by the European Union – NextGenerationEU.

## Data availability statement

The RNA-seq and ATAC-seq datasets generated during this study are publicly available in the Gene Expression Omnibus repository under accession number GSE332970 and GSE332971.

## Author contributions

Xingyu Fan: Formal analysis, Investigation, Methodology, Visualization, Writing – original draft, Writing – review & editing

Bram Torenvliet: Formal analysis, Investigation, Methodology, Software, Visualization, Writing – original draft, Writing – review & editing

Alexandros Galaras: Formal analysis, Investigation, Methodology, Software, Visualization, Writing – review & editing

Tanvir Hossain: Writing – review & editing

Lincon Hasda: Investigation, Writing – review & editing

Martin E van Royen: Formal analysis, Investigation, Writing – review & editing Helmuth Gehart: Writing – review & editing

Lili Zhao: Writing – review & editing

Eleni Katsoni-Resources, Writing – review & editing

Tsung Wai Kan: Resources, Writing – review & editing

Panagiotis Moulos: Writing – review & editing

Shringar Rao: Writing – review & editing

Farzin Pourfarzad: Writing – review & editing

Javier Frias Aldeguer: Investigation, Writing – review & editing

Sylvia F. Boj: Funding acquisition, Supervision, Writing – review & editing

Pantelis Hatzis: Methodology, Writing – review & editing

Robert-Jan Palstra: Conceptualization, Formal analysis, Funding acquisition, Investigation, Methodology, Supervision, Visualization, Writing – original draft, Writing – review & editing

Tokameh Mahmoudi: Conceptualization, Funding acquisition, Methodology, Supervision, Writing – original draft, Writing – review & editing

