## Supplementary figures and images for "Hepatitis B virus protein X promotes hepatocyte plasticity and survival in a differentiated human liver organoid system"

### Supplementary Fig. 1

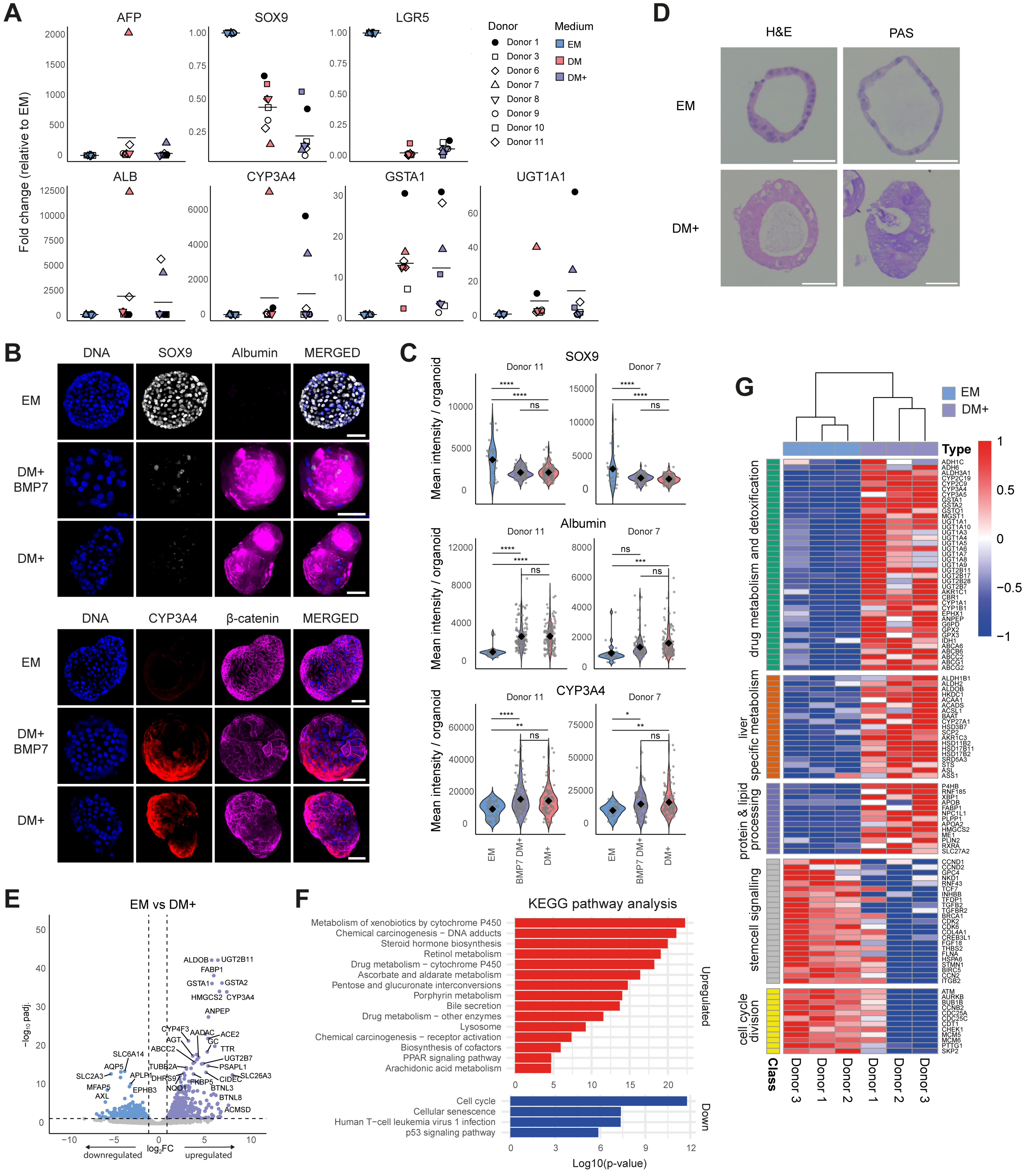

### Supplementary Fig. 2

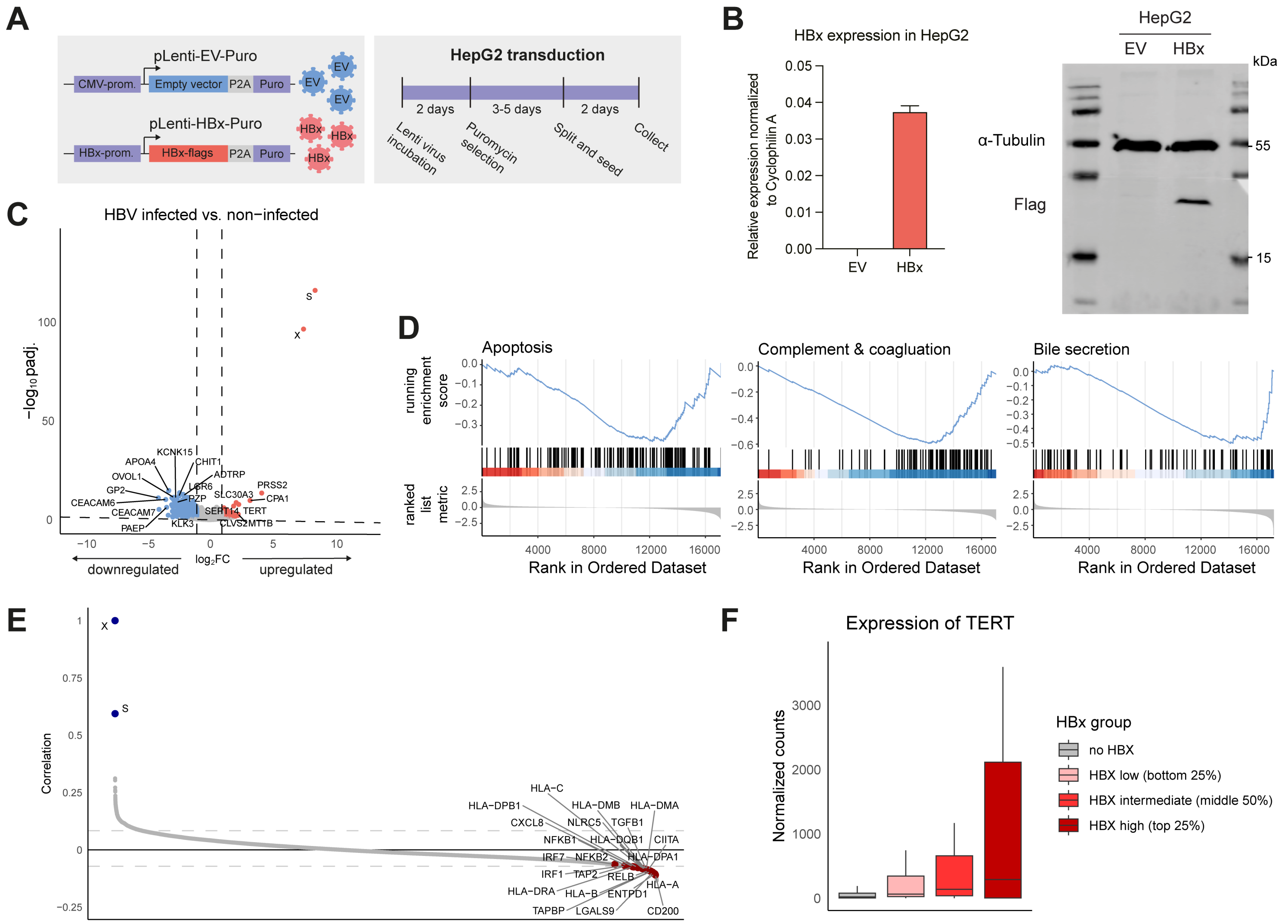

### Supplementary Fig. 3

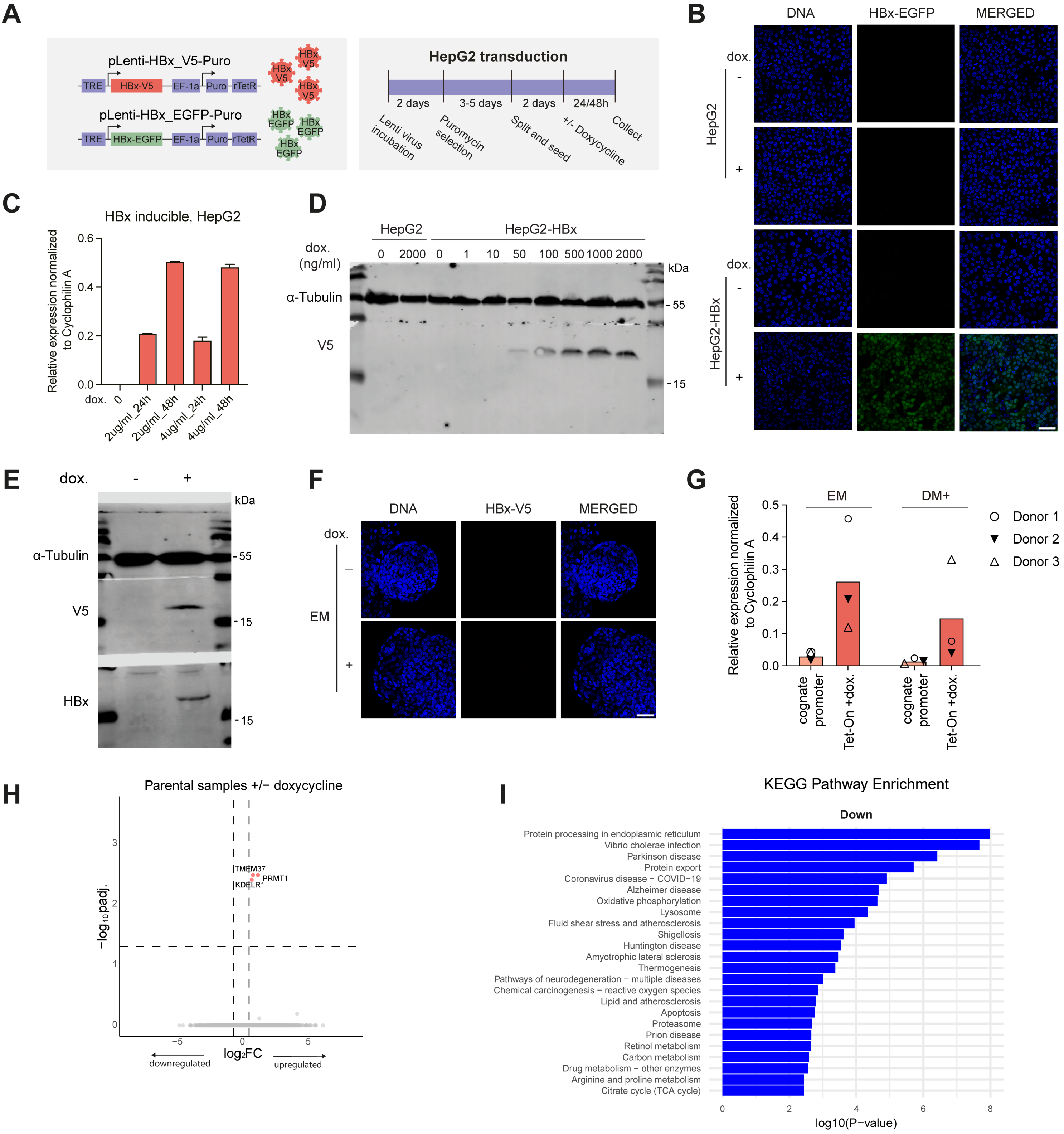

### Supplementary Fig. 4

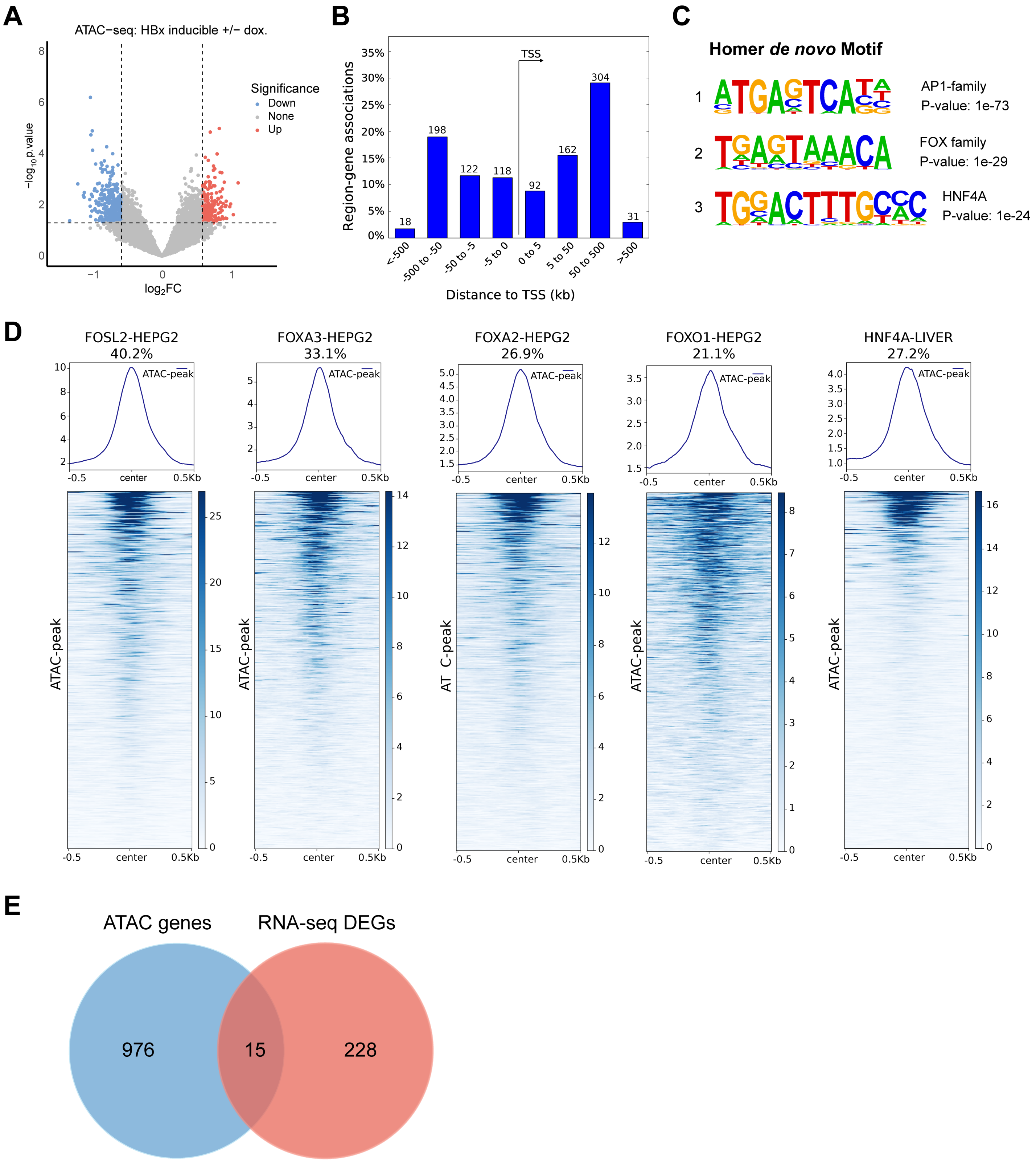

### Supplementary Fig. 5

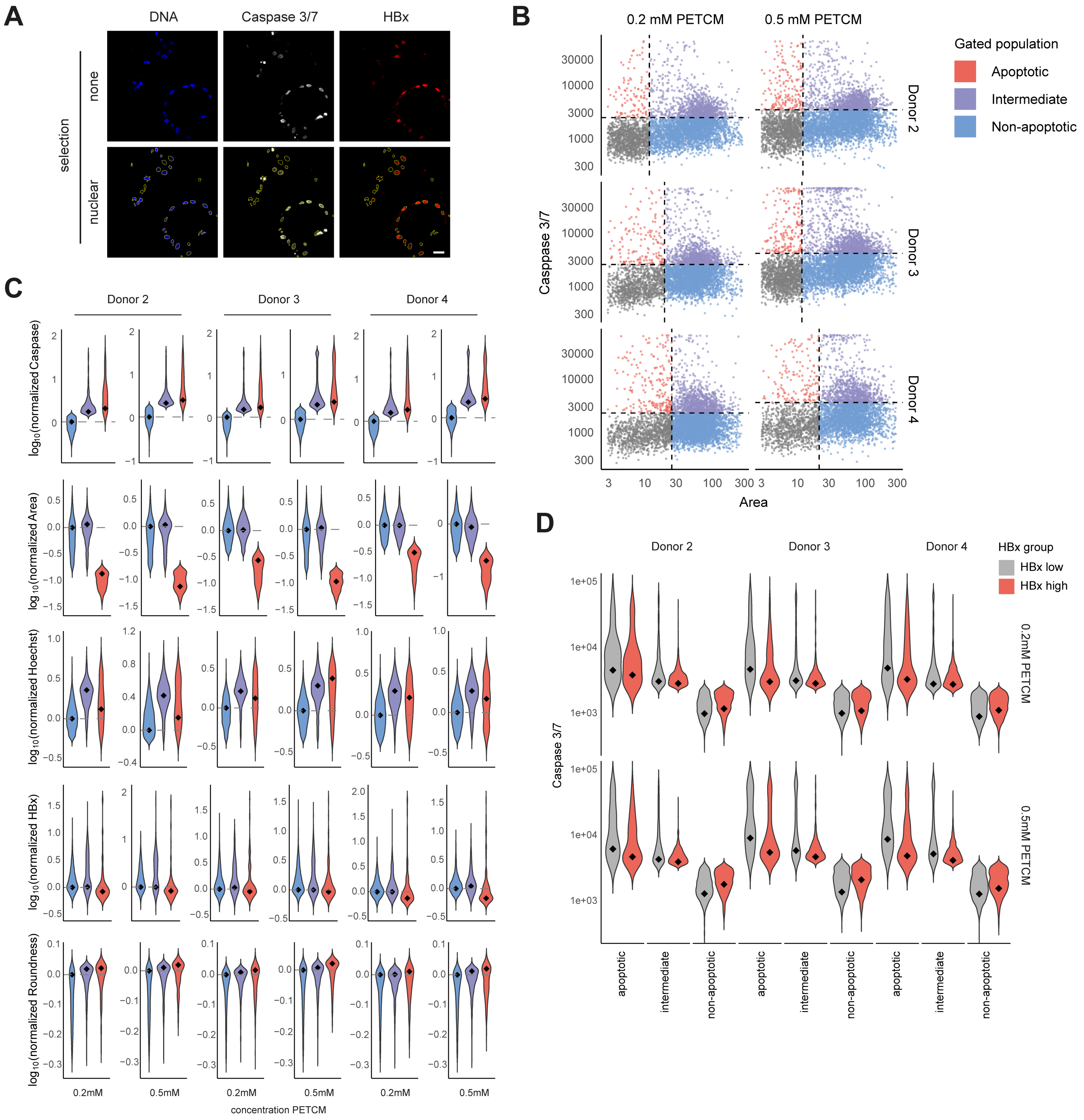

### Supplementary Fig. 6

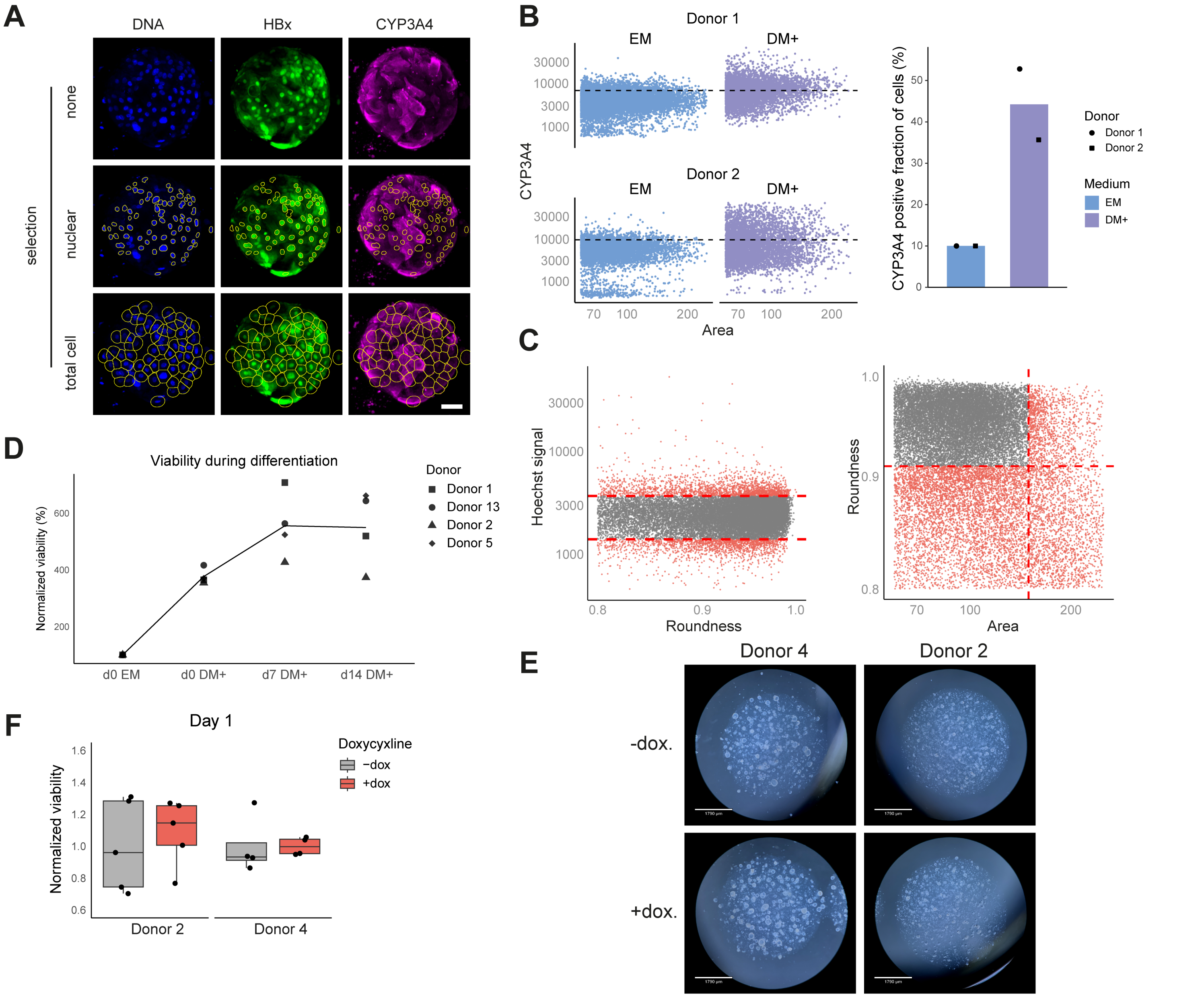
