## Supplementary Information for "Hepatitis B virus protein X promotes hepatocyte plasticity and survival in a differentiated human liver organoid system"

**Title page**

**Title**

[#] These authors contributed equally.

* Corresponding author

Corresponding author

Tokameh Mahmoudi

Address: Departments of Pathology and Urology, Erasmus University Medical Center, Netherlands

**Table of content**

Title page 1

Title 1

Authors 1

Affiliations 1

Corresponding author 2

Table of content 3

Materials and Methods 5

Establishment and maintenance of liver organoids 5

Differentiation of liver organoids 7

Cell culture 8

Generation of lentivectors 8

Lentiviral particle generation 9

Lentiviral transduction and induction of Tet-on system 9

RNA isolation and Reverse Transcription-quantitative PCR (RT-qPCR) 11

RNA sequencing 11

RNA-sequencing data processing and analysis 13

Analysis of public microarray data 14

TCGA RNA-seq. re-analysis against the HBV genome 14

Pathway and gene set enrichment analyses 15

Assay of transposase-accessible chromatin with sequencing (ATAC-seq.) 16

Western blotting 17

Immunofluorescence staining and fluorescence microscopy 18

High throughput imaging analysis 19

Ex vivo drug treatment 20

Colony formation and survival 20

Statistics 21

Additional information microscopy (HUB) 21

Additional information microscopy (Erasmus Medical Center) 22

Supplemental Figure legends 22

Supplemental Figure 1. 22

Supplemental Figure 2. 23

Supplemental Figure 3. 24

Supplemental Figure 4. 25

Supplemental Figure 5. 26

Supplemental Figure 6. 27

References for Materials and Methods 29

#### Materials and Methods

##### Establishment and maintenance of liver organoids

Patient material and patient-derived organoids were collected at two institutions under approved ethical protocols and after obtaining informed consent from all patients. At HUB Organoids B.V. (HUB), collection and use of material were approved by the Medical Ethics Review Committee and the Biobank Research Ethics Committee (TcBio) under the HUB-Cancer protocol (12-093) and TcBio release protocol #24-237. Six of the organoid lines are available through the Foundation Hubrecht Organoid Biobank (https://www.hubrechtorganoidbiobank.org/). Donor-derived material at Erasmus Medical Center was used for research as described previously (1).

Organoids were derived from histologically normal liver tissue obtained from patients undergoing partial hepatectomy for tumour resection, or from liver tissue of healthy donors, and biobanked as previously described (1-4). Briefly, liver tissues were first washed by Dulbecco's Modified Eagle Medium (DMEM, Sigma) supplemented with 1 % foetal calf serum (FCS, Sigma) and 0.1 % penicillin/streptomycin (PS, Sigma), minced with scalpels and enzymatically dissociated to single cells by incubation with Collagenase D (Roche). The resulting cell suspension was filtered through a 70 μm cell strainer (Corning), washed with Ad+++ medium (Advanced DMEM/F12 (Gibco) supplemented with 1% PS, 10 mM HEPES (Gibco) and 1% GlutaMax (Gibco)), and resuspend in basement membrane extract (BME) solution consisting of 2/3 BME type 2 (R&D systems) and 1/3 Ad+++. Cells embedded in BME were seeded in suspension plates (Greiner) and incubated at 37°C for 30 minutes to allow gelation. To initiate organoid formation, wells were overlaid with Isolation medium (IM) consisting of Ad+++ supplemented with 1X B27 supplement minus vitamin A (Gibco), 1.25 mM N-acetyl-L-cysteine (Sigma), 10 mM Nicotinamide (Sigma), 10 nM recombinant human (Leu15)-gastrin I (Tocris), 50 ng/mL recombinant human epidermal growth factor (hEGF, Peprotech), 100 ng/mL recombinant human fibroblast growth factor 10 (hFGF10, Peprotech), 25 ng/mL recombinant human hepatocyte growth factor (hHGF, Peprotech), 10 µM Forskolin (Sigma), 5 µM A83-01 (Tocris), 25 ng/mL Noggin (Peprotech), 10 µM Y27632 Rho Kinase (ROCK) Inhibitor (Sigma), and either 250 ng/mL Rspo-3 (R&D systems) or a combination of 1X N2 supplement (Gibco), 20% (vol/vol) Rspo-1 conditioned medium (prepared in-house) and 1.25% (vol/vol) Wnt3a conditioned medium (prepared in-house). Cultures were incubated at 37 °C in a humidified atmosphere containing 5% $CO_2$.

Once organoids were established, IM was replaced with Expansion Medium (EM) consisting of Ad+++ supplemented with 1X B27 supplement minus vitamin A, 1.25 mM N-acetyl-L-cysteine, 10 mM Nicotinamide, 10 nM recombinant human (Leu15)-gastrin I, 50 ng/mL recombinant human EGF, 100 ng/mL recombinant human FGF10, 25 ng/mL recombinant human HGF, 10 µM Forskolin, 5 µM A83-01, and either 250 ng/mL Rspo-3 or a combination of 1X N2 supplement, 20% (vol/vol) Rspo-1 conditioned medium and 1.25% (vol/vol) Wnt3a conditioned medium. The medium was refreshed 3 times per week.

Liver organoids were passaged at a 1:4 to 1:8 ratio every 7–10 days depending on organoid density and size. To passage the organoids, the BME solution containing the organoids was collected by adding 500 µg/mL Dispase II (Sigma) for 1 hour (h) and washed once with Ad+++. The organoid pellet was then resuspended in 100 ul TrypLE Express (Gibco) dissociated into organoid fragments by mechanical pipetting. After removing TrypLE Express by washing with Ad+++, the pellet was resuspended in fresh BME solution (2/3 BME and 1/3 Ad+++) and plated, overlaid with EM supplemented with 10 μM ROCK inhibitor for enhanced cell survival. The ROCK inhibitor was removed after 2–3 days of recovery in EM. For cryopreservation, dissociated organoids were resuspended in Recover™ Cell Culture Freezing Medium (Gibco), transferred to CryoTube™ vials (Thermo Scientific), and frozen at -80 °C for 2–3 days in a freezing container before being transferred to liquid nitrogen for long-term storage. Subsequent establishment of organoid cultures from frozen stocks was performed as follows; Cryopreserved liver organoid samples were rapidly thawed in a 37 °C water bath, resuspended in 10 mL of Ad+++ and centrifuged at 200 × g for 5 min at room temperature. The organoid pellet was then resuspended in BME solution, allowed to solidify in a 37 °C incubator, and overlaid with EM supplemented with 10 μM ROCK Inhibitor. ROCK inhibitor was omitted from EM after the first medium change. Organoids were routinely tested for mycoplasma contamination and authenticated by single nucleotide polymorphism (SNP) profiling to confirm sample identity.

Differentiation of liver organoids

Differentiation of liver organoids was started once organoids reached an average diameter between 100 and 250 μm, typically 3-7 days post-passaging. In order to facilitate differentiation, EM was removed from the culture wells, and replaced with differentiation medium (DM; Ad+++ supplemented with 1X B27 supplement minus vitamin A, 1 mM N-acetylcysteine, 10 nM recombinant human [Leu15]-gastrin I, 50 ng/mL recombinant human hEGF, 25 ng/mL recombinant human hHGF, 0.5 μM A83-01, 10 μM N-[N-(3,5-Difluorophenacetyl)-L-alanyl]-S-phenylglycine t-butyl ester (DAPT, Sigma), 3 μM Dexamethasone (Sigma), 25 ng/mL BMP7 (Peprotech), 100 ng/mL recombinant human FGF19 (hFGF19, Peprotech)) with or without 1X N2 supplement, or Improved Differentiation Medium (DM+; World Intellectual Property Organization Patent WO2017149025A1), consisting of DM with the addition of 100 μM

Carbamyolcholine Chlorine (Sigma), 3 μM CHIR 99021 (Tocris), 25 μM iCRT3 (Sigma) and 3 μM Stem MACS IWP-2 (Miltenyi Biotec).

Cell culture

HepG2 and HEK293T cells were obtained from ATCC. HepG2.2.15 cells were obtained from CCTCC. The identity of the cells was authenticated by the manufacturers by short tandem repeats profiling and cell lines were routinely tested for mycoplasma. HepG2, HepG2.2.15 and HEK293T cells were cultured in DMEM supplemented with 10% FCS and 1% P/S, and incubated at 37 °C in a humidified atmosphere containing 5% $CO_2$. For the maintenance of HepG2.2.15 cells, 380 mg/L G418 (Invitrogen) was added into the medium for 2 weeks after thawing from the biobank to ensure selection of positively transduced cells.

Generation of lentivectors

A pLenti-C-Myc-DDK-P2A-Puro Lentiviral Gene Expression Vector (pLenti-EV-Puro, Origene) was used to generate a plasmid containing the HBx coding sequence driven by the cognate HBx promoter (pLenti-HBx-Puro) via molecular cloning. The DNA fragment containing the HBx promoter and HBx coding sequence tagged with two FLAG epitopes (2×FLAG) was amplified from HepG2.2.15 cDNA by Polymerase Chain Reaction (PCR) (primer pair listed in Supplementary Materials). Both the empty vector pLenti-EV-Puro and the PCR-amplified fragment were digested with the XbaI (New England Biolabs) and XhoI (New England Biolabs) restriction enzymes at 37 °C overnight, followed by ligation using a T4 DNA ligase kit (Promega) according to the manufacturer’s instructions.

For the construction of a Tet-on inducible lentiviral vector expressing HBx (pLenti-HBx_V5_Puro), the HBx coding sequence tagged with V5 epitopes was amplified by PCR from the pLenti-HBx-Puro described above (primer pair listed in Supplementary Materials). The PCR product and the TLCV2, pLentiCRISPR v2 all-in-one doxycycline-inducible vector (Addgene), were digested with the AgeI-HF (New England Biolabs) and NheI-HF (New England Biolabs) restriction enzymes and ligated using a T4 DNA ligase kit according to the manufacturer's instructions. The Tet-on inducible HBx-EGFP fusion vector (pLenti-HBx_EGFP_Puro) was generated from the resulting pLenti-HBx_V5_Puro plasmid. First, the EGFP sequence was amplified by PCR from the dCas13d-dsRBD-APEX2 vector (Addgene). Next, this EGFP sequence was C-terminally fused to the HBx sequence inside the HBx_V5_Puro plasmid via restriction enzyme digesting by AscI (New England Biolabs) and NheI-HF, followed by ligation using a T4 DNA ligase kit according to the manufacturer's instructions. Correct plasmid assembly was verified by whole plasmid sequencing prior to downstream analyses.

Lentiviral particle generation

For lentiviral particle generation, 7–8 × $10^6$ HEK293T cells were seeded in 10 cm dishes 8–12 h prior to transfection. Cells were co-transfected with 6 µg lentiviral expression plasmid, 2 µg VSV-G envelope plasmid (pMD2.G, Addgene), and 4.5 µg second-generation packaging plasmid (psPAX2, Addgene). Plasmids were diluted in 500 µL serum-free DMEM and combined with 500 µL serum-free DMEM containing 125 µL of 1 mg/mL polyethyleneimine (Sigma). After incubation for 30 min at room temperature, the transfection mixture was diluted with 9 mL serum-free DMEM and added to the cells. After 6–8 h, the medium was replaced with Ad+++ for lentiviral particle collection. Viral supernatants were collected at 24 and 48 h post-transfection, filtered through a 0.45 µm cellulose acetate membrane (Millipore), and stored at −80 °C until further usage.

Lentiviral transduction and induction of Tet-on system

For transduction, 1 × $10^5$ HepG2 cells were seeded per well in a 12-well plate and cultured overnight. The following day, medium was replaced with 250 µL lentiviral supernatant supplemented with 750 µL DMEM supplemented with 10% FCS and 1% PS (DMEM++). Cells were incubated for 48 h, after which the medium was replaced with fresh culture medium DMEM++ and cells were allowed to recover for an additional 48 h. Transduced cells were selected using 2 µg/mL puromycin (Invitrogen) for 5-7 days.

For lentiviral transduction of liver organoids, organoids from four wells of a 24-well plate were harvested, washed with Ad+++, and dissociated to an approximately 80% single cell suspension using TrypLE Express as described previously. The pellet was resuspended in 1 mL lentiviral supernatant and distributed across four wells of a 48-well plate. Plates were sealed using parafilm and centrifuged at 600 × g, 32 °C for 1 h. Following centrifugation, cells were resuspended and incubated at 37 °C in a humidified atmosphere containing 5% $CO_2$ for 5–6 h. Cells were then collected, washed with Ad+++, embedded in BME, and overlaid with EM supplemented with 10 µM ROCK inhibitor and 25 ng/mL Noggin. After 5–7 days, ROCK inhibitor and Noggin were omitted from the medium, and selection was initiated using 2 µg/mL puromycin for 5–7 days. Resistant organoids were passaged and allowed to recover for an additional 5–7 days prior to downstream applications. For induction of HBx expression in Tet-on organoids, 2 µg/mL doxycycline (Sigma) was added to either EM or DM+ for 24 h.

RNA isolation and Reverse Transcription-quantitative PCR (RT-qPCR)

Cell or organoid pellets from 2–3 wells of a 24-well plate were collected, washed, and lysed in 1 mL of TRI Reagent (Sigma) for RNA isolation. Chloroform (200 µL per 1 mL TRI Reagent) was added, and samples were vigorously shaken for 15 seconds followed by centrifugation at 12,000 × g for 15 min at 4 °C. The aqueous phase was collected, and RNA was precipitated by centrifugation at 12,000 × g for 15 min at 4 °C. The RNA pellet was washed twice with 1 mL 75% ethanol and centrifuged at 7,500 × g for 5 min at 4 °C. Pellets were briefly air-dried and resuspended in 30–50 µL nuclease-free water.

For downstream applications, 300–1000 ng RNA was treated with DNase I (Invitrogen) to remove residual genomic DNA, followed by cDNA synthesis using a High-Capacity cDNA Reverse Transcription Kit (Applied Biosystems) according to the manufacturer's instructions. cDNA was diluted 1:5–1:10 in nuclease-free water prior to quantitative PCR.

RT–qPCR was performed with SYBR-based detection (GoTaq qPCR Master Mix (Promega)) on a CFX Connect Real-Time PCR Detection System (BioRad). Each reaction contained 4 µL cDNA, 1 µL (10 µM) primer mix, and 5 µL master mix. Amplification was carried out with an initial denaturation at 95 °C for 3 min, followed by 40 cycles of 95 °C for 10 s and 60 °C for 30 s. Product specificity was confirmed by melting curve analysis, and no-RT controls were routinely included. Primer sequences are listed in Supplementary Materials.

RNA sequencing

RNA samples were quantitated and quality-controlled on a Qubit 4 Fluorometer (Invitrogen) with the Qubit RNA HS Assay Kit (Invitrogen) and on an Agilent

TapeStation 4150 system with the High Sensitivity RNA ScreenTape assay (Agilent), respectively, according to the manufacturers' instructions.

Libraries were prepared using the QuantSeq 3′ mRNA-Seq Library Prep Kit FWD with Unique Dual Indices (Lexogen), following the manufacturer's instructions. Briefly, up to 500 ng of RNA was used for first-strand synthesis, followed by RNA template removal. Second-strand synthesis was initiated using random primers containing compatible linker sequences at their 5′ end. In-line barcodes were introduced during second-strand synthesis, followed by magnetic bead-based purification, amplification and subsequent purification.

Library quality and quantity were assessed using the Agilent TapeStation 4150 with the High Sensitivity D1000 assay kit and the Qubit 4 Fluorometer with the Qubit dsDNA HS Assay Kit, respectively. Libraries were quantified, pooled equimolarly, and used for Adapter Conversion PCR (AC-PCR) amplification using the Universal Library Conversion Kit (App-A), Version 1.0 (MGI Tech Co., Ltd.), according to manufacturer's instructions.

The quality of the purified AC-PCR product was evaluated using the Agilent TapeStation 4150 and the High Sensitivity D1000 assay kit, while quantity was assessed using the Qubit 4 Fluorometer with the Qubit dsDNA HS Assay Kit. This was followed by DNA denaturation, single-strand circularization, enzymatic digestion, enzymatic digestion product cleanup, and quality control, according to manufacturer's instructions.

Finally, the ssCirDNA was used for DNB preparation and single-end sequencing on the DNBSEQ-G400 platform at the BSRC Alexander Fleming Genomics Facility, using the G400 App-A FCS SE100 High-throughput Sequencing Set (MGI Tech Co., Ltd.), according to manufacturer's instructions.

Raw FASTQ files were processed using Trim Galore! to remove low-quality bases from read ends and terminal ambiguous nucleotides (N bases). Reads shorter than 50 bp after trimming were discarded. The remaining reads were aligned against the human reference genome build hg38 using a two-step method. First, reads were mapped to the reference transcriptome with HISAT2 (5), and the remaining unmapped reads were aligned with Bowtie2 (6). Read counting on the 3' UTRs was conducted using the metaseqR2 bioconductor package (7) with default settings.

RNA-sequencing data processing and analysis

All RNA-sequencing data analysis was performed in RStudio. First, lowly expressed genes (fewer than 10 counts in more than two-third of samples) were filtered out from the read counts table. Next counts were normalized for inherent systematic or experimental biases using the Bioconductor package DESeq2 (8) by variance stabilizing transformation. Paired differential gene expression analysis followed by DESeq2. Differential expression was assessed using the Wald test, and p-values were adjusted for multiple testing using the Benjamini-Hochberg method. Significance thresholds for (adjusted) p-values and log2 fold change were not fixed but adapted to each biological comparison. Specifically, different thresholds were applied depending on the expected magnitude of transcriptional changes, such as comparisons involving gene overexpression versus differentiation states.

Analysis of public microarray data

Publicly available microarray data were obtained from Gene Expression Omnibus (accession: GSE63859). The dataset was downloaded using the GEOquery R package (9). As the data were already normalized, no additional normalization was performed. Probe annotations were mapped to gene symbols, and probes without annotation were removed. Samples were filtered to retain relevant experimental groups based on metadata. For genes represented by multiple probes, expression values were averaged to obtain a single value per gene.

TCGA RNA-seq. re-analysis against the HBV genome

BAM files corresponding to 371 patients from The Cancer Genome Atlas liver hepatocellular carcinoma (LIHC) cohort were obtained through the Genomic Data Commons (GDC) portal following controlled-access approval. For the cases where multiple samples were available, only one sample was retained based on the following priority: I) primary tumour samples, II) solid tissue normal samples, and III) recurrent tumour samples.

Sequencing reads were first aligned to a custom reference genome comprising 9311 HBV sequences corresponding to 8 genotypes (multi-genotype) obtained from HBVdb v.62.0 (10) using STAR (11) with chimeric alignment detection (--chimSegmentMin 15, --chimJunctionOverhangMin 12, --chimMainSegmentMultNmax 1) enabled to identify candidate host–viral junctions. The remaining unmapped reads were re-aligned to the HBV-only multi-genotype reference using Bowtie2 (6), with the parameters --very-sensitive-local --dovetail, to improve sensitivity and recover viral reads not captured during the initial alignment. Only reads with mapping quality (MAPQ) ≥20 were retained for further analysis.

Counting of reads was performed with FeatureCounts (subread-2.0.6). Genes were retained if they had a minimum count of 10 in at least 25% of samples. Then, samples were classified based on HBV infection status. Specifically, cases were considered HBV-infected if they were clinically annotated as HBV-positive and/or exhibited evidence of viral transcription (combined normalized count greater than 5 across the X and S genes). All remaining samples were classified as HBV-non-infected.

Differential expression analysis was performed with metaseqR2 (7) using the DESeq2 algorithm (8) between HBV infected and non-infected samples. As differentially expressed we considered genes with FDR < 0.05 and absolute log2 fold change > 0.58.

Pathway and gene set enrichment analyses

Pathway and gene set-based analyses were performed using complementary approaches to assess biological processes associated with gene expression changes. For over-representation analysis, differentially expressed genes were mapped to KEGG pathway annotations (12) after conversion from gene symbols to Entrez identifiers using the org.Hs.eg.db annotation database (13). Enrichment analysis was conducted using the clusterProfiler package (14), and the most significantly enriched pathways for each comparison were selected for visualization.

To estimate pathway activity at the sample level, gene set variation analysis (GSVA) was performed using the GSVA package (14), normalized expression data and predefined gene sets. Enrichment scores were calculated in a non-parametric manner for each sample.

In addition, gene set enrichment analysis (GSEA) was conducted using ranked gene lists to evaluate the enrichment of selected biological processes. Gene sets were derived from KEGG pathway annotations, and for broader biological themes, multiple related pathways were combined into unified gene sets prior to analysis. All enrichment analyses were performed using functions implemented in the clusterProfiler package (15).

Assay of transposase-accessible chromatin with sequencing (ATAC-seq.)

ATAC-seq libraries were prepared using the ATAC-Seq Kit (Active motif) according to the accompanying protocol. Library quality was assessed using the Agilent TapeStation 4150 system with the High Sensitivity D1000 assay kit, while library DNA quantity was measured using the Qubit 4 Fluorometer (Invitrogen) with the Qubit dsDNA HS Assay Kit. Libraries were pooled equimolarly and used for Adapter Conversion PCR (AC-PCR) amplification using the Universal Library Conversion Kit (App-A), Version 1.0 (MGI Tech Co., Ltd.), according to manufacturer's instructions.

The quality of the purified AC-PCR product was evaluated using the Agilent TapeStation 4150 and the High Sensitivity D1000 assay kit, while quantity was assessed using the Qubit 4 Fluorometer with the Qubit dsDNA HS Assay Kit. This was followed by DNA denaturation, single-strand circularization, enzymatic digestion, enzymatic digestion product cleanup, and quality control, according to manufacturer's instructions.

Finally, the ssCirDNA was used for DNB preparation and sequencing on the DNBSEQ-G400 platform at the "BSRC Alexander Fleming" Genomics Facility, using the App-D-converted G400 FCL PE100 High-throughput Sequencing Set (MGI Tech Co., Ltd.), according to manufacturer's instructions.

Raw FASTQ files were trimmed to remove Nextera adapter sequences and aligned to the human reference genome (hg38) using Bowtie2 (6). Duplicate reads and reads mapped to extra chromosomes were removed from the resulting BAM files prior to peak calling. Peaks were identified using MACS2 (16) using the parameters --broad --nomodel --shift 100 --extsize 200. Read counting and differential accessibility analysis were performed using the bioconductor package DiffBind (17) with the EDGER algorithm (18). Peaks overlapping ENCODE blacklisted regions were excluded from downstream analyses. Read counts were quantified for each peak and normalized using edgeR. Only peaks with a minimum of 10 read counts across samples were retained for further analysis. Regions with an absolute log2 fold change greater than 0.58 and a p-value < 0.05 were defined as differentially accessible regions (DARs). Peak annotation was performed using annotatePeaks.pl from HOMER (Hypergeometric Optimization of Motif EnRichment) (18) with default parameters. Motif enrichment analysis was subsequently conducted using HOMER.

ChIP-seq data obtained in HEPG2 cells or Human liver for potential motif bound transcription factors was downloaded from the ENCODE portal (www.encodeproject.org) (19). To investigate the overlap between HBx DAR ATAC-peaks and transcription factors we used the tool ‘Intersect the intervals of two datasets’ tool in Galaxy with default settings (20). Heatmaps were generated using the ‘computeMatrix’ (reference-point and centre of region output options) and ‘plotHeatmap’ tools in Galaxy.

Western blotting

Cells or organoid pellets were lysed in NP-40 immunoprecipitation buffer containing 1% NP-40 (Thermo Scientific), 25 mM Tris-HCl (pH 7.4; Cytiva), 150 mM NaCl (Sigma), 1 mM EDTA (Sigma), 5% glycerol (Sigma), 1 U/mL EDTA-free protease inhibitor cocktail (Roche), and 1 mM DTT (Sigma). Lysates were incubated on ice for 30 min and clarified by centrifugation at 14,000 rpm for 10 min at 4 °C. Supernatants were collected, mixed with 1× Laemmli loading buffer (Thermo Scientific), and boiled for 5 min at 95 °C. Proteins were resolved by 15% SDS-PAGE and subsequently transferred onto a PVDF membrane. Following transfer, the membrane was blocked with 5% non-fat dry milk in PBST (0.1% Tween20 in PBS) for 1 h at room temperature. The membranes were then incubated with primary antibodies overnight at 4°C, followed by a 2 h incubation with corresponding secondary antibodies at room temperature. Protein bands were visualized and imaged using the Odyssey CLx Imaging System (LI-COR Biosciences). Antibodies used are listed in Supplementary Methods.

Immunofluorescence staining and fluorescence microscopy

Organoids were collected in Eppendorf tubes, washed with PBS, and fixed in 4% paraformaldehyde (PFA) in PBS for 20 min at room temperature. Following fixation, organoids were washed three times with PBS and pelleted by centrifugation at 100 × g for 2 min at room temperature. For permeabilization and blocking, pellets were incubated in PBS containing 0.3% Triton X-100 and 5% bovine serum albumin (BSA) for 1 h at room temperature. Organoids were washed three times with washing buffer (0.5% BSA in PBS) and incubated overnight at 4 °C with primary antibodies (listed in Supplementary Materials) diluted in antibody incubation buffer (2.5% BSA in PBS).

The following day, organoids were washed three times with washing buffer and incubated with fluorophore-conjugated secondary antibodies (1:1000) in antibody incubation buffer for 2 h at room temperature in the dark. After three washes, DNA was stained with 5 µg/mL Hoechst 33342 (Molecular Probes) for 30 min at room temperature. Organoids were then washed three times in PBS and either placed in mounting medium (Dakocytomation) and covered with coverslips or resuspended in PBS and transferred to 96-well clear-bottom plates (Revvity) for imaging.

Confocal imaging was performed using a Leica Stellaris 5 Laser Scanning Microscope, an Opera Phenix Plus High-Content Screening System (Revvity) or an ImageXpress Confocal HT.ai, high-content imaging system (Molecular Devices). During imaging by the ImageXpress or Opera Phenix high-content imaging systems, an initial pre-scanning was conducted, at either an 4x or 10x magnification to identify field of view (FOV) containing organoid material, based on the brightfield or Hoechst channel. The identified FOVs were subsequently imaged at 20x or 40x magnification. Images were acquired as z-stacks. For organoid-level quantification, maximum intensity projections were used, whereas single-cell analyses were performed on individual z-planes.

Representative images were processed using Fiji (ImageJ). Brightness and contrast were adjusted using linear transformations applied uniformly across the full image. Identical adjustments were applied to all images within each comparison.

##### High throughput imaging analysis

Downstream high-content image analysis was performed using the integrated INCarta (Molecular Devices) and Harmony (Revvity) software, for the ImageXpress or Opera Phenix microscope, respectively. All quantifications include more than 65 individual organoids per condition. Organoid-level quantification of mean fluorescence intensity (MFI) relied on a brightfield-based segmentation of organoid outline, followed by a quantification of MFI within the region of interest (ROI). Single cell-level quantifications were performed by segmentation of nuclear outline, using Hoechst signal. The cytoplasmic region was approximated by applying a 3 µm or 10 µm isotropic expansion to the nuclear segmentation mask. Next, MFI across all staining channels was quantified for both the nuclear, and cytoplasmic ROI. Image analysis data was output from the INCarta and Harmony software and subsequently analysed and visualized using Rstudio. Prior to analysis, debris and artifacts were excluded from the dataset by on object area and Hoechst intensity. Modes were estimated from kernel density functions (Gaussian kernel) by identifying the density maximum. Summary statistics were calculated as arithmetic means unless stated otherwise. Data visualization was performed using ggplot2 (21).

Ex vivo drug treatment

pLenti-HBx_V5_Puro transduced liver organoids were split at a 1:1 ratio and filtered through 20-100 µm strainers prior to seeding. After 4-6 days in EM, organoids were differentiated in DM+ for 10 days. Following the differentiation, 2 µg/mL doxycycline was added to the medium for 24 h before apoptosis induction by PETCM (MedChemExpress) for 12 h. Sample preparation for high-content image analysis was performed as described above. Caspase 3/7 activity was visualized using CellEvent™ Caspase-3/7 Detection Reagents Green (ThermoFisher), which was incubated (1:500) for 60 min at 37 °C, before fixation. Samples were imaged and analysed as described above.

Colony formation and survival

pLenti-HBx_EGFP_Puro and pLenti-HBx_V5_Puro transduced organoids were cultured in a 96-well Flat Clear Bottom Black Polystyrene TC-treated Microplates (Corning) for 5-7 days in EM followed by a 10 day differentiation in DM+. Transgene expression was induced by addition of 2 µg/mL dox. for 24 h, after which organoids were treated with 0.05 mM PETCM for up to 8 days. In order to evaluate organoid growth and survival, organoids were imaged every 24 h during PETCM treatment at 5x magnification. For each well, a single FOV was captured at the centre of the BME dome, with 8 imaging planes captured 80 µm apart, centred around this location.

Organoid outline was determined by manual segmentation. Cell viability was evaluated using CellTiter-Glo® 3D Cell Viability Assay (Promega) according to the manufacturer's instructions. Each experimental condition was performed with at least two biological replicates.

#### Statistics

Statistical analyses were performed in R. Data are presented as mean ± SD unless otherwise indicated. Statistical significance was assessed using one-way ANOVA followed by Tukey's multiple comparisons test or using paired Wilcoxon signed-rank testing. P values < 0.05 were considered statistically significant. Statistical significance is indicated as follows: *P < 0.05, **P < 0.01, ***P < 0.001, ****P < 0.0001; ns, not significant.

#### Additional information microscopy (HUB)

Make and model of microscope: ImageXpress Confocal HT.ai, high-content imaging system (Molecular Devices)

Type, magnification, and numerical aperture of the objective lenses: 4x (NA 0.2, dry), 20x (NA 0.95, water immersion) & 40x (NA 1.15, water immersion)

Temperature: room temperature

Imaging medium: PBS

Camera make and model: CMOS camera with a 2048 × 2048 image sensor format (6.5 × 6.5 μm pixel size) and a peak quantum efficiency of 82%.

Acquisition software: MetaXpress (Molecular Devices)

Additional information microscopy (Erasmus Medical Center)

Make and model of microscope: Opera Phenix High Content Screening system equipped with a Yokogawa microlens enhanced wide view Nipkow spinning disk (Revvity).
Type, magnification, and numerical aperture of the objective lenses: 5x (NA 0.16, dry; For Colony formation and survival), 20x (NA 0.4, dry; For Immunofluorescence/ fluorescence imaging) & 40x (NA 1.1, water immersion; For Immunofluorescence/ fluorescence imaging)
Temperature: room temperature (Immunofluorescence/ fluorescence imaging), 37 °C (Colony formation and survival)
Imaging medium: PBS
Camera make and model: 16bit sCMOS camera with a 2160 x 2160 image sensor format and a peak quantum efficiency of 82%
Acquisition software: Harmony version 5.2 (Revvity)

#### Supplemental Figure legends

##### Supplemental Figure 1.

DM+ condition improves hepatocyte-like organoid differentiation. (A) Gene expression analysis of progenitor and hepatocyte marker genes of HLOs in different culture media by RT-qPCR; n = 8 biological replicates. Expression is displayed as fold change, normalized to expression in EM. Horizontal bar indicates the mean fold change across biological replicates. (B) Representative immunofluorescence staining of organoids prior to and following differentiation, with or without exposure to BMP7 prior to differentiation. Donor 11, scale bar; 50 µm. (C) Violin plots of mean fluorescent intensity (MFI) per organoid, per medium. Grey dots represent the MFI determined in an individual organoid and black diamonds represent the mean MFI across all organoids in the corresponding condition. Statistical significance was assessed using one-way ANOVA followed by Tukey's multiple comparisons test. (D) Representative H&E and PAS staining of organoids prior to and following differentiation. Donor 11, scale bar; 50 μm. (E) Volcano plot of differential gene expression between organoids in EM and DM+. Significantly upregulated and downregulated genes were defined by adjusted p value < 0.05 and $|\log_2$ fold change$| > 1$ and are highlighted. (F) KEGG pathway enrichment analysis performed on differentially expressed genes identified between organoids in EM and DM+. (G) Heatmap of normalized gene expression for genes associated with progenitor or hepatocyte cell identities across organoids in EM and DM+.

**Supplemental Figure 2.**

Cognate promoter driven HBx expression in HepG2 cells and HBV infected patient cohort analysis. (A) Left, schematic overview of lentiviral construct design. A lentiviral plasmid was generated containing the cognate HBx promoter and the HBx sequence, fused to two flag tags (pLenti-HBx-Puro). An empty vector plasmid (pLenti-EV-Puro) was used as a control. Right, schematic representation of the experimental outline. HepG2 cells were incubated with lentivirus for 2 days, followed by a selection in 1 μg/mL puromycin for 3–5 days. Positively transduced cells were split and seeded for 2 days prior to collection. (B) Left, gene expression analysis of HBx in empty vector and HBx-expressing HepG2 cells by RT-qPCR; n = 3 technical replicates. Right, Protein expression of HBx in empty vector and pLenti-HBx-Puro transduced HepG2 cells by Western-Blot. (C) Volcano plot of differential gene expression between HBV infected and non-infected patient. Significantly upregulated and downregulated genes were defined by adjusted p value < 0.05 and $|\log_2$ fold change$| > 1$ and are highlighted. (D)

Gene set enrichment analysis plot showing negative enrichment of apoptosis, and liver-function associated genes in HBV infected patients. (E) Correlation plot of gene expression with HBx levels across patient samples. Genes are ranked by correlation coefficient, with the top and bottom 5% correlation coefficient values indicated by grey horizontal dotted lines. Viral genes (X and S) are highlighted in dark blue, and immune response–associated genes in dark red. (F) TERT expression stratified by HBx levels. Patient samples were grouped into HBx-negative, low (bottom 25%), intermediate (middle 50%), and high (top 25%) expression, and TERT expression is shown as box plots for each group.

**Supplemental Figure 3.**

Inducible Tet-On system driven HBx expression in HepG2 cells and HLOs. (A) Left, schematic overview of lentiviral construct design. A lentiviral plasmid was created containing the HBx sequence fused with either a V5-tag (pLenti-HBX_V5_Puro) or an EGFP sequence (pLenti_HBx_EGFP-Puro), following a Tet-on TRE promoter. Right, schematic representation of the experimental outline. HepG2 cells were incubated with lentivirus for 2 days, followed by a selection in 1 µg/mL puromycin for 3–5 days. Positively transduced cells were split and seeded for 3 days prior to collection. During the final day of culture, 2 µg/mL dox. was added for 24 h to induce the overexpression of HBx. (B) Representative fluorescence staining of HBx-EGFP transduced and wild-type (wt) HepG2 cells ± dox. Scale bar; 50 µm. (C) Gene expression analysis of HBx in HepG2 (pLenti-HBX_V5_Puro) cells cultured ± dox. at varying doxycycline concentrations, by qPCR; n = 3 technical replicates. (D) Protein expression of HBx-V5 in transduced and wt HepG2 cells ± dox. at varying dox. concentrations, by Western blotting. (E) Raw western blotting image showing HBx expression in (pLenti-HBx_V5_Puro) transduced organoids cultured in DM+, with alpha (α) Tubulin functioning as the housekeeping control, donor 2, corresponding to Fig. 2C. (F) Representative immunofluorescence staining of parental organoids ± dox. Scale bar; 50 µm. (G) Gene expression analysis of HBx in EM- and DM+-HLOs (transduced by pLenti-HBx-Puro or pLenti-HBX_V5_Puro) by qPCR; n = 3 biological replicates. pLenti-HBX_V5_Puro transduced HLOs were cultured + 2 µg/mL dox. for 24 h prior to collection. (H) Volcano plot of differential gene expression between parental organoids ± 2 µg/mL dox.. Significantly upregulated and downregulated genes were defined by adjusted p value < 0.05 and $|\log_2$ fold change$|$ > 1 and are highlighted. (I) KEGG pathway enrichment analysis performed on differentially expressed genes identified between pLenti-HBX_V5_Puro transduced HLOs ± dox.

**Supplemental Figure 4.**

HBx-induced changes in chromatin accessibility and transcription factor recruitment. (A) Volcano plot of differential gene accessibility between pLenti-HBX_V5_Puro transduced DM+-HLOs ± dox, n = 2 biological replicates. Significant DARs were defined by p value < 0.05 and $|\log_2$ fold change$|$ > 0.58 and are highlighted. (B) Bar plot of frequency distribution of DAR distance to the nearest TSS determined by GREAT analysis, n = 2 biological replicates. (C) Density plots and occupancy analysis using ENCODE ChIP-seq data show significant overlap of DARs with FOSL2, FOXA3, FOXA2, and FOXO1 in HEPG2 cells, and HNF4A in liver tissue samples. (D) Top enriched motifs of the DARs identified by Homer analysis between pLenti-HBX_V5_Puro transduced DM+-HLOs ± dox, n = 2 biological replicates. (E) Venn diagram of the intersection between genes associated with DARs and differentially expressed genes (both up- and down-regulated) found in RNA-seq. analysis of pLenti-HBX_V5_Puro transduced DM+-HLOs ± dox., identifying 15 common genes.

**Supplemental Figure 5.**

Exogenous expression of HBx in DM+-HLOs attenuates the induction of apoptosis. (A) Representative immunofluorescence staining of DM+-HLOs +dox. treated with 0.5 mM PETCM, highlighting segmentation of the nuclear outline in yellow. Nuclear outline was determined using DNA signal. Donor 3, scale bar; 25 µm. (B) Scatterplots of mean activated caspase 3/7 intensity against area of individual nuclei in PETCM-treated DM+-HLOs. Vertical and horizontal dotted lines indicate the gating used to classify nuclei as either non-apoptotic, intermediate or apoptotic. Gating was performed per donor-treatment combination, with area and caspase 3/7 thresholds set to the lowest and highest quartiles, respectively. (C) Violin plots of normalized caspase, area, Hoechst, HBx, and nuclear roundness across different cell populations. Data are shown per donor–treatment combination. Violin fill colours indicate cell populations, with blue, purple, and red corresponding to non-apoptotic, intermediate, and apoptotic cells, respectively. Normalization was performed per donor–treatment combination relative to the non-apoptotic population. The mode of each distribution is indicated by a black diamond and was determined from the peak of the kernel density distribution. Horizontal dotted lines mark the mode of the non-apoptotic population. (D) Violin plots of nuclear caspase 3/7 intensity in PETCM treated cells, across the different cell populations. Nuclei from each population were classified as HBx-high or HBx-low, based on whether they displayed HBx intensity values in the top or bottom 50% of the population.

**Supplemental Figure 6.**

Exogenous expression of HBx results in dedifferentiation of DM+-HLOs and evasion of apoptosis. (A) Representative immunofluorescence images of +dox. treated DM+-

HLOs, highlighting segmentation of the nuclear and total cellular outline in yellow. Nuclear outline was determined using DNA signal. Total cellular outline was estimated by performing a 10 µm radial expansion of the nuclear outline. Donor 12, scale bar; 50 µm. (B) Left, scatterplot of mean CYP3A4 intensity within the total cellular outline, against nuclear area of individual cells, EM- and DM+-HLOs. Horizontal dotted line indicated threshold applied for CYP3A4-positivity, set at the 90th percentile of EM-HLO cell intensity, per donor. Right, barplot indicating CYP3A4-positive percentage of the cell population within EM and DM+. Height of the barplots display the mean percentage across included donors, per culture medium. (C) Left, scatterplot of nuclear roundness against nuclear Hoechst signal. Right, scatterplot of nuclear area against nuclear roundness. In both plots, dotted red lines indicate gating applied to the corresponding parameter. Only cells containing nuclei within the gates, displayed in grey, were included in downstream analysis, while cells shown in light-red were excluded from downstream analysis. (D) Normalized viability of organoids at different timepoints during differentiation. Black line indicates the average viability across donors. Each individual data point represents the average of $n \geq 3$ technical replicates. Viability was normalized to the viability at time of organoid seeding (d0 EM) and determined using Cell-titer Glo 3D. (E) Representative brightfield images showing organoid density before PETCM treatment. Scale bar; 1790 µm. (F) Boxplot showing normalized viability of organoids at day 1 of PETCM treatment. Viability was normalized to the viability of the -dox. condition and determined using Cell-titer Glo 3D.

### Refences:


1. De Crignis E, Hossain T, Romal S, Carofiglio F, Moulos P, Khalid MM, et al. Application of human liver organoids as a patient-derived primary model for HBV infection and related hepatocellular carcinoma. Elife. 2021;10.
2. Romal S, Hossain T, Mahmoudi T. Generation, Maintenance and HBV Infection of Human Liver Organoids. Bio Protoc. 2022;12(6):e4358.
3. Hossain T, Romal S, Mahmoudi T. Production of Recombinant Hepatitis B virus (HBV) and Detection of HBV in Infected Human Liver Organoids. Bio Protoc. 2022;12(8):e4392.
4. Rao S, Romal S, Torenvliet B, Slotman JA, Huijs T, Mahmoudi T. A 3D organoid platform that supports liver-stage P.falciparum infection can be used to identify intrahepatic antimalarial drugs. Heliyon. 2024;10(10):e30740.
5. Kim D, Paggi JM, Park C, Bennett C, Salzberg SL. Graph-based genome alignment and genotyping with HISAT2 and HISAT-genotype. Nat Biotechnol. 2019;37(8):907-15.
6. Langmead B, Salzberg SL. Fast gapped-read alignment with Bowtie 2. Nat Methods. 2012;9(4):357-9.
7. Fanidis D, Moulos P. Integrative, normalization-insusceptible statistical analysis of RNA-Seq data, with improved differential expression and unbiased downstream functional analysis. Brief Bioinform. 2021;22(3).
8. Love MI, Huber W, Anders S. Moderated estimation of fold change and dispersion for RNA-seq data with DESeq2. Genome Biol. 2014;15(12):550.
9. Davis S, Meltzer PS. GEOquery: a bridge between the Gene Expression Omnibus (GEO) and BioConductor. Bioinformatics. 2007;23(14):1846-7.
10. Hayer J, Jadeau F, Deleage G, Kay A, Zoulim F, Combet C. HBVdb: a knowledge database for Hepatitis B Virus. Nucleic Acids Res. 2013;41(Database issue):D566-70.
11. Dobin A, Davis CA, Schlesinger F, Drenkow J, Zaleski C, Jha S, et al. STAR: ultrafast universal RNA-seq aligner. Bioinformatics. 2013;29(1):15-21.
12. Kanehisa M, Goto S. KEGG: kyoto encyclopedia of genes and genomes. Nucleic Acids Res. 2000;28(1):27-30.
13. org.Hs.eg.db: genome wide annotation for Human [
14. Hanzelmann S, Castelo R, Guinney J. GSVA: gene set variation analysis for microarray and RNA-seq data. BMC Bioinformatics. 2013;14:7.
15. Yu G. Thirteen years of clusterProfiler. Innovation (Camb). 2024;5(6):100722.
16. Zhang Y, Liu T, Meyer CA, Eeckhoute J, Johnson DS, Bernstein BE, et al. Model-based analysis of ChIP-Seq (MACS). Genome Biol. 2008;9(9):R137.
17. Stark R BG. DiffBind: differential binding analysis of ChIP-Seq peak data. 2011 [Available from: https://bioconductor.org/packages/release/bioc/vignettes/DiffBind/inst/doc/DiffBind.pdf. .
18. Heinz S, Benner C, Spann N, Bertolino E, Lin YC, Laslo P, et al. Simple combinations of lineage-determining transcription factors prime cis-regulatory elements required for macrophage and B cell identities. Mol Cell. 2010;38(4):576-89.
19. Sloan CA, Chan ET, Davidson JM, Malladi VS, Strattan JS, Hitz BC, et al. ENCODE data at the ENCODE portal. Nucleic Acids Res. 2016;44(D1):D726-32.
20. Galaxy C. The Galaxy platform for accessible, reproducible, and collaborative data analyses: 2024 update. Nucleic Acids Res. 2024;52(W1):W83-W94.
21. ggplot2: Elegant Graphics for Data Analysis [
