## Supplementary Materials for "Hepatitis B virus protein X promotes hepatocyte plasticity and survival in a differentiated human liver organoid system"

### 1 Antibodies

| Name | Citation | Supplier | Cat no. | Clone no. |
|---|---|---|---|---|
| ANTI-FLAG M2 antibody (mouse monoclonal) | RRID:AB_259529 | Sigma-Aldrich | F3165 | M2 |
| Anti-TUBA4A (TUBA1) Antibody | RRID:AB_477579 | Sigma-Aldrich | T5168 | B-5-1-2 |
| Hepatitis B Virus X Monoclonal Antibody | RRID:AB_325418 | Invitrogen | MA1-081 | X36C |
| V5 Tag Monoclonal Antibody | RRID: AB_2556564 | Invitrogen | R960-25 | SV5-Pk1 |
| Human Albumin Polyclonal Antibody | RRID:AB_67018 | Bethyl Laboratories | A80-229A | N/A |
| Anti-SOX9 antibody | RRID:AB_2728660 | Abcam | ab185966 | EPR14335-78 |
| Anti-human CYP3A4 | RRID: AB_2876366 | BioIVT | PAP 011 | N/A |
| Purified Anti-β-Catenin (mouse monoclonal) | RRID:AB_397555 | BD Transduction Laboratories | 610154 | 14 |

### 2 Cell lines

| Name | Citation | Supplier | Cat no. | Passage no. | Authentication test method |
|---|---|---|---|---|---|
| HepG2 | RRID:CVCL_0027 | ATCC | HB-8065 | 6-8 | STR |
| HepG2.2.15 | RRID:CVCL_L855 | CCTCC | CCTCC-GDC0141 | 6-8 | STR |
| HEK 293T | RRID:CVCL_0063 | ATCC | CRL-3216 | 6-8 | STR |

### 3 Primers

| Name | Sequence | Supplier |
|---|---|---|
| PCR primer (PLenti-HBx-Puro) | Forward: TCGCATCATGCTCTAGAGGCCTATTGATTGGAAAG TATG Reverse: GGACTTACACTCGAGCTTATCGTCGTCATCCTTGT AATCCTTATCGTCGTCATCCTTGTAATCGCCTCCG GCAGAGGTGAAAAAGTT | Integrated DNA Technologies |
| PCR primer (PLenti-HBx_V5_Puro) | Forward: TCGCATCATGCACCGGTATGGCTGCTAGGCT GTGCTG Reverse: GGACTTACAGCTAGCCGCGCGCTTAGCCTCC CGTAGAATCGAGACCGAGGAGAGGGTTAGGG ATAGGCTTACCGCCTCCCTTATCGTCGTCATC CTTGTAATCCTTATCGTCGTCATCCTTGTAATC GAATTCGGGCGCGCCGCCTCCGGCAGAGGTG AAAAAGTTGCA | Integrated DNA Technologies |
| PCR primer (PLenti-HBx_EGFP_Puro) | Forward: TCGCATCATGCGCGGCGCGCCTAAGATGGTGAG CAAGGGCGAG Reverse: TTAGCGCTAGCGCGCTTAGCGCTCCTTGTACAGC TCGTCCATGCC | Integrated DNA Technologies |
| RT_qPCR primer Human AFP | Forward: CTTTGGGCTGCTCGCTATGA Reverse: GCATGTTGATTTAACAAGCTGCT | Integrated DNA Technologies |
| RT_qPCR primer Human SOX9 | Forward: AGCGAACGCACATCAAGAC Reverse: CTGTAGGCGATCTGTTGGGG | Integrated DNA Technologies |

| | | |
|---|---|---|
| RT_qPCR primer Human<br>LGR5 | Forward:<br>AGGTCTGGTGTGTTGCTGAGG<br>Reverse:<br>TGAAGACGCTGAGGTTGGAAGG | Integrated DNA Technologies |
| RT_qPCR primer Human<br>ALB | Forward:<br>TGCAACTCTTCGTGAAACCTATG<br>Reverse:<br>ACATCAACCTCTGGTCTCACC | Integrated DNA Technologies |
| RT_qPCR primer Human<br>CYP3A4 | Forward:<br>TGTGCCTGAGAACACCAGAG<br>Reverse:<br>GTGGTGGAAATAGTCCCGTG | Integrated DNA Technologies |
| RT_qPCR primer Human<br>GSTA1 | Forward:<br>CTGCCCGTATGTCCACCTG<br>Reverse:<br>AGCTCCTCGACGTAGTAGAGA | Integrated DNA Technologies |
| RT_qPCR primer Human<br>UGT1A1 | Forward:<br>CATGCTGGGAAGATACTGTTGAT<br>Reverse:<br>GCCCGAGACTAACAAAAGACTCT | Integrated DNA Technologies |
| RT_qPCR primer Human<br>HBx | Forward:<br>GGCATACTTCAAAGACTGTTTGTTT<br>Reverse:<br>CGCAGACCAATTTATGCCTAC | Integrated DNA Technologies |
| RT_qPCR primer Human<br>Cyclophilin A | Forward:<br>TCATCTGCACTGCCAAGACTG<br>Reverse:<br>CATGCCTTCTTTCACTTTGCC | Integrated DNA Technologies |
| RT_qPCR primer Human<br>18s | Forward:<br>AAACGGCTACCACATCCAAG<br>Reverse: | Integrated DNA Technologies |

| | |
|---|---|
| | CCTCCAATGGATCCTCGTTA |

### 4 Biological samples

| Description | Source | Identifier |
|---|---|---|
| Liver organoid Donor 1, established from healthy liver tissue | Department of Transplantation | Erasmus University Medical Center |
| Liver organoid Donor 2, established from healthy liver tissue | Department of Transplantation | Erasmus University Medical Center |
| Liver organoid Donor 3, established from normal liver tissue | Universitair Medisch Centrum Utrecht | HUB Organoids B.V. (HUB) |
| Liver organoid Donor 4, established from healthy liver tissue | Department of Transplantation | Erasmus University Medical Center |
| Liver organoid Donor 5, established from healthy liver tissue | Department of Transplantation | Erasmus University Medical Center |
| Liver organoid Donor 6, established from normal liver tissue | Universitair Medisch Centrum Utrecht | HUB Organoids B.V. (HUB) |
| Liver organoid Donor 7, established from normal liver tissue | Universitair Medisch Centrum Utrecht | HUB Organoids B.V. (HUB) |
| Liver organoid Donor 8, established from normal liver tissue | Universitair Medisch Centrum Utrecht | HUB Organoids B.V. (HUB) |
| Liver organoid Donor 9, established from normal liver tissue | Universitair Medisch Centrum Utrecht | HUB Organoids B.V. (HUB) |
| Liver organoid Donor 10, established from normal liver tissue | Universitair Medisch Centrum Utrecht | HUB Organoids B.V. (HUB) |
| Liver organoid Donor 11, established from normal liver tissue | Universitair Medisch Centrum Utrecht | HUB Organoids B.V. (HUB) |
| Liver organoid Donor 12, established from healthy liver tissue | Department of Transplantation | Erasmus University Medical Center |

| | | |
|---|---|---|
| Liver organoid Donor 13, established from healthy liver tissue | Department of Transplantation | Erasmus University Medical Center |

### 5 Software

| Software name | Manufacturer | Version |
|---|---|---|
| GraphPad Prism 10 | GraphPad Software | N/A |
| RStudio | RStudio | 4.5.1 |
| DESeq2 | Bioconductor package | 1.48.1 |
| GEOquery R package | Bioconductor package | 2.76.0 |
| org.Hs.eg.db annotation database | Bioconductor package | 3.21.0 |
| clusterProfiler package | Bioconductor package | 4.16.0 |
| GSVA package | Bioconductor package | 2.2.0 |
| Ggplot2 package | Hadley Wickham | 3.5.2 |
| Fiji (Image J) | NIH | 1.54p |
| Adobe illustrator | Adobe | 29.8.6 |
| Harmony | Revvity | 5.2 |
| Odyssey CLx Imaging System | LI-COR Biosciences | 5.2 |
| MetaXpress | Molecular Devices | 6.7.2.290 |
| INCarta | Molecular Devices | 2.8.0912.228 |

### 6 Other (e.g. drugs, proteins, vectors etc.)

| | | |
|---|---|---|
| DMEM | Gibco | Cat# 41966052 |
| FCS | Gibco | Cat# F7524 |
| PS | Sigma | Cat# P4333 |
| Collagenase D | Roche | Cat# 11088858001 |
| 70 μm cell strainer | Corning | Cat# 352350 |
| Advanced DMEM/F12 | Gibco | Cat# 12634010 |
| HEPES | Gibco | Cat# 15630080 |
| GlutaMax | Gibco | Cat# 35050061 |

| | | |
|---|---|---|
| BME | R&D systems | Cat# 3533-010-02 |
| Suspension plates | Greiner | Cat# 662102 |
| 1X B27 supplement minus vitamin A | Gibco | Cat# 12587001 |
| N2 supplement | Gibco | Cat# 17502048 |
| N-acetyl-L-cysteine | Sigma | Cat# A9165 |
| Rspo-3 | R&D Systems | Cat# 3500-RS/CF |
| Nicotinamide | Sigma | Cat# N0636 |
| recombinant human (Leu15)-gastrin I | Tocris | Cat# 3006 |
| hEGF | Peprotech | Cat# AF-100-15 |
| hHGF | Peprotech | Cat# 100-3 |
| hFGF10 | Peprotech | Cat# 100-26 |
| Forskolin | Sigma | Cat# F3917 |
| A83-01 | Tocris | Cat# 2939 |
| Noggin | Peprotech | Cat# 120-10C |
| ROCK Inhibitor | Sigma | Cat# SCM075 |
| DispaseII | Sigma | Cat# D4693 |
| TryplE Express | Gibco | Cat# 12604013 |
| Recover™ Cell Culture Freezing Medium | Gibco | Cat# 12648010 |
| CryoTube vials | Thermo Scientific | Cat# 5000-1012 |
| DAPT | Sigma | Cat# D5942 |
| Dexamethasone | Sigma | Cat# D4902 |
| BMP7 | Peprotech | Cat# 120-03 |
| hFGF19 | Peprotech | Cat# 100-32 |
| Carbamyolcholine Chlorine | Sigma | Cat# C4382 |
| CHIR 99021 | Tocris | Cat# 4423 |
| iCRT3 | Sigma | Cat# SML0211 |
| Stem MACS IWP-2 | Miltenyi Biotec | Cat# 130-105-335 |
| G418 | Invitrogen | Cat# ant-gn-1 |

| | | |
|---|---|---|
| pLenti-C-Myc-DDK-P2A-Puro Lentiviral Gene Expression Vector | Origene | Cat# PS100092 |
| XbaI | New England Biolabs | Cat# R0145S |
| XhoI | New England Biolabs | Cat# R0146S |
| AgeI-HF | New England Biolabs | Cat# R3552S |
| NheI-HF | New England Biolabs | Cat# R3131S |
| AscI | New England Biolabs | Cat# R0558S |
| T4 DNA ligase kit | Promega | Cat# M1801 |
| TLCV2 | Addgene | Cat# 87360 |
| dCas13d-dsRBD-APEX2 | Addgene | Cat# 154939 |
| pMD2.G | Addgene | Cat# 12259 |
| psPAX2 | Addgene | Cat# 12260 |
| PEI | Polyscience | Cat# 23966 |
| 0.45 μm cellulose acetate membrane | Millipore | Cat# SLHAR33SS |
| Puromycin | Invitrogen | Cat# ANT-PR-1 |
| Doxycycline | Sigma | Cat# D9891 |
| TRI Reagent | Sigma | Cat# T3809 |
| DNase I | Invitrogen | Cat# 18068-015 |
| High-Capacity cDNA Reverse Transcription Kit | Applied Biosystems | Cat# 4368813 |
| GoTaq qPCR Master Mix kit | Promega | Cat# A6001 |
| ATAC-Seq Kit | Active Motif | Cat# 53150 |
| NP-40 | Thermo Scientific | Cat# 85124 |
| Tris | Cytiva | Cat# GE17-1321-01 |
| NaCl | Sigma | Cat# S9888 |
| EDTA | Sigma | Cat# E9884 |
| Glycerol | Sigma | Cat# G7893 |
| EDTA-free protease inhibitor cocktail | Roche | Cat# 11836170001 |
| DTT | Sigma | Cat# 43816 |

| | | |
|---|---|---|
| Laemmli loading buffer | Thermo Scientific | Cat# J60015.AD |
| PVDF membrane | Millipore | Cat# IPVH00010 |
| TWEEN20 | Sigma | Cat# P1379 |
| 680RD donkey anti-mouse | LICORbio | Cat# 926-68072 |
| 680RD donkey anti-rabbit | LICORbio | Cat# 926-68073 |
| BSA | Sigma | Cat# A3294 |
| Hoechst 33342 | Molecular Probes | Cat# 33258 |
| Alexa Fluor 488 donkey anti-mouse IgG (H+L) | Invitrogen | Cat# A-21202 |
| Alexa FluorÂ 594 Donkey Anti-Mouse IgG (H+L) | Invitrogen | Cat# A-21203 |
| Alexa Fluor 647 donkey anti-mouse IgG (H+L) | Invitrogen | Cat# A-31571 |
| Alexa Fluor_ 594 donkey anti-goat IgG (H+L) | Invitrogen | Cat# A-11058 |
| Alexa Fluor 647 donkey anti-Rabbit IgG (H+L) | Invitrogen | Cat# A-31573 |
| Mounting Medium | Dakocytomation | Cat# S3023 |
| Corning® 96-well Flat Clear Bottom Black Polystyrene TC-treated Microplates | Corning | Cat# 3603 |
| ViewPlate-96 Black, Optically Clear Bottom plate | Revvity | Cat# 6005225 |
| CellEvent™ Caspase-3/7 Detection Reagents Green | Invitrogen | Cat# C10423 |
| Triton X-100 | Sigma | Cat# X100RS |
| CellTiter-Glo® 3D Cell Viability Assay | Promega | Cat# G9681 |
| PETCM | MedchemExpress | Cat# HY-103349 |
| PFA | Electron Microscopy | Cat# 15710 |
